# Correction of amblyopia in cats and mice after the critical period

**DOI:** 10.1101/2021.05.03.442423

**Authors:** Ming-fai Fong, Kevin R. Duffy, Madison P. Leet, Christian T. Candler, Mark F. Bear

## Abstract

Monocular deprivation early in development causes amblyopia, a severe visual impairment. Prognosis is poor if therapy is initiated after an early critical period. However, clinical observations have shown that recovery from amblyopia can occur later in life when the non-deprived (fellow) eye is removed. The traditional interpretation of this finding is that vision is improved by relieving interocular suppression in primary visual cortex. However, an alternative explanation is that elimination of activity in the fellow eye establishes conditions in visual cortex that enable the weak connections from the amblyopic eye to gain strength. Here we show in cats and mice that temporary inactivation of the fellow eye is sufficient to promote a full and enduring recovery from amblyopia at ages when conventional treatments fail. Thus, connections serving the amblyopic eye are capable of substantial plasticity beyond the critical period, and this potential is unleashed by reversibly silencing the fellow eye.

## Introduction

Amblyopia is a prevalent form of visual disability that emerges during infancy when inputs to the visual cortex from the two eyes are poorly balanced (Simons, 2005). The most common causes of amblyopia are strabismus and asymmetric refraction, but the most severe form—deprivation amblyopia—arises from opacities or obstructions of vision (e.g., by cataract). The current standard of care is to restore clarity (e.g. by cataract extraction) and focus, and then promote recovery of the weak amblyopic eye by temporarily patching the fellow eye (Wallace et al., 2018). However, the effectiveness of occlusion therapy is limited by poor compliance, variable recovery outcomes, and a significant risk of recurrence. Additionally, occlusion therapy is largely ineffective if it is initiated after age 10 (DeSantis, 2014) or, in the case of deprivation amblyopia, after the first year of life (Birch & Stager, 1996). The need for improved treatments for amblyopia is widely acknowledged (Falcone, Hunter, & Gaier, 2021; Elizabeth M Quinlan & Lukasiewicz, 2018).

Studies over many decades in cats and monkeys have shown how temporary monocular deprivation (MD) sets in motion a series of changes in primary visual cortex (V1) that degrade vision through the deprived eye. As in humans, these changes can be reversed by temporarily occluding the fellow eye, but the effectiveness of this procedure is again limited to a brief critical period (Blakemore, Garey, & Vital-Durand, 1978; Blakemore & Van Sluyters, 1974). Rodents have become the dominant animal model for study of the synaptic basis of amblyopia, and an important recent development has been the finding from multiple laboratories that diverse manipulations, all culminating in reduced inhibition by a population of cortical interneurons, can allow recovery from the effects of MD at ages beyond the classically defined critical period (Bavelier, Levi, Li, Dan, & Hensch, 2010; Hensch & Quinlan, 2018; Sengpiel, 2014). As exciting as these results are, however, most of the manipulations used in mice and rats to promote adult recovery have limited applicability to humans. Some are not feasible in a therapeutic setting (e.g., interneuron transplantation (Davis et al., 2015)) and others require systemic exposure to agents that have actions beyond visual cortex (e.g., cholinesterase inhibitors (Morishita, Miwa, Heintz, & Hensch, 2010) and ketamine (Grieco et al., 2020)), and none have demonstrated clinical efficacy in a human study to date (e.g., fluoxetine (Huttunen et al., 2018), citalopram (Lagas, Black, Russell, Kydd, & Thompson, 2019), levodopa (Repka et al., 2015), donepezil (Chung, Li, Silver, & Levi, 2017)). Indeed, some treatments that work robustly in rodents have yielded disappointing results when investigated in other species with a more differentiated visual system, like the cat (e.g., (Holman, Duffy, & Mitchell, 2018; Vorobyov, Kwok, Fawcett, & Sengpiel, 2013)). It is now recognized that studies limited to a single species, particularly rats or mice with primitive visual systems, may be unreliable guides to human amblyopia treatment (D. Mitchell & Sengpiel, 2018).

In the current study we took a “bedside to bench” perspective. One interesting observation in the human clinical literature is that significant recovery from amblyopia can sometimes occur in adults when the normal (fellow) eye is damaged or removed (enucleated) following injury or disease (El Mallah, Chakravarthy, & Hart, 2000; Kaarniranta & Kontkanen, 2003; Klaeger-Manzanell, Hoyt, & Good, 1994; Rahi et al., 2002; Vereecken & Brabant, 1984). Similar findings in cat and monkey models have been interpreted to mean that a mechanism contributing to amblyopia is the continuous suppression of responses to the deprived eye by activity through the fellow eye (Harwerth, Smith, Crawford, & von Noorden, 1984; Hendrickson, Boles, & McLean, 1977; Hoffmann & Lippert, 1982; Kratz & Spear, 1976). However, an alternative hypothesis is that eliminating activity in the strong eye via enucleation causes a homeostatic adjustment in the properties of synaptic plasticity that allows experience through the amblyopic eye to drive synaptic strengthening (Cho & Bear, 2010; Cooper & Bear, 2012). If this idea is correct, merely silencing the fellow eye temporarily should be sufficient to enable a lasting recovery. Thus, the “suppression removal” hypothesis predicts that amblyopia will be immediately mitigated by silencing the fellow eye, but only for as long as that eye is inactive, whereas the “synaptic strengthening” hypothesis predicts that recovery will emerge gradually and outlast the period of retinal inactivation. We have conducted experiments in two species, mouse and cat, to distinguish among these alternatives. Our results show that temporary inactivation of the fellow eye after long-term MD can permanently restore vision through both eyes.

## Results

### Temporary monocular inactivation potentiates non-inactivated eye responses in mouse V1

We first sought to characterize the impact of temporary retinal inactivation on the cortical responses to visual stimulation in neurotypical mice, initiated at postnatal day (P) 47 which is after the critical period has ended (Gordon & Stryker, 1996). Recording electrodes were implanted into V1 layer 4 to measure baseline visual evoked potentials (VEPs) in awake, head-fixed animals (**Figs. 1A-B, S1**). In agreement with previous studies, under baseline conditions the response to stimulation of the contralateral eye was approximately double that of the ipsilateral eye at baseline (**Figs. 1C-D**). The contralateral retina was then inactivated for 1-2 days using the voltage-gated sodium channel blocker tetrodotoxin (TTX), injected into the vitreous humor. During the period of inactivation, responses to stimulation of the inactivated contralateral eye predictably fell to noise levels (**Fig. 1C**). Meanwhile, responses to stimulation of the non-inactivated ipsilateral eye rose dramatically (**Fig. 1D**). We continued to track contralateral and ipsilateral VEPs for several days as the effect of TTX dissipated and retinal activity returned. Even one week later, non-inactivated eye responses remained significantly elevated above baseline (**Figs. 1D-E**). Importantly, responses measured during stimulation of the contralateral eye returned to baseline levels (**Figs. 1C, E**), indicating that there is no lasting decrement in inactivated eye responsiveness. These results demonstrate that removal of the influence of the dominant eye immediately augments responses through the non-dominant eye, consistent with interocular suppression. In addition, the data suggest that a brief period of monocular retinal inactivation also sets the stage for lasting potentiation of responses to the non-inactivated eye. These responses remain potentiated well beyond the period of inactivation and without long-term consequences to the inactivated eye.

**Figure 1:**
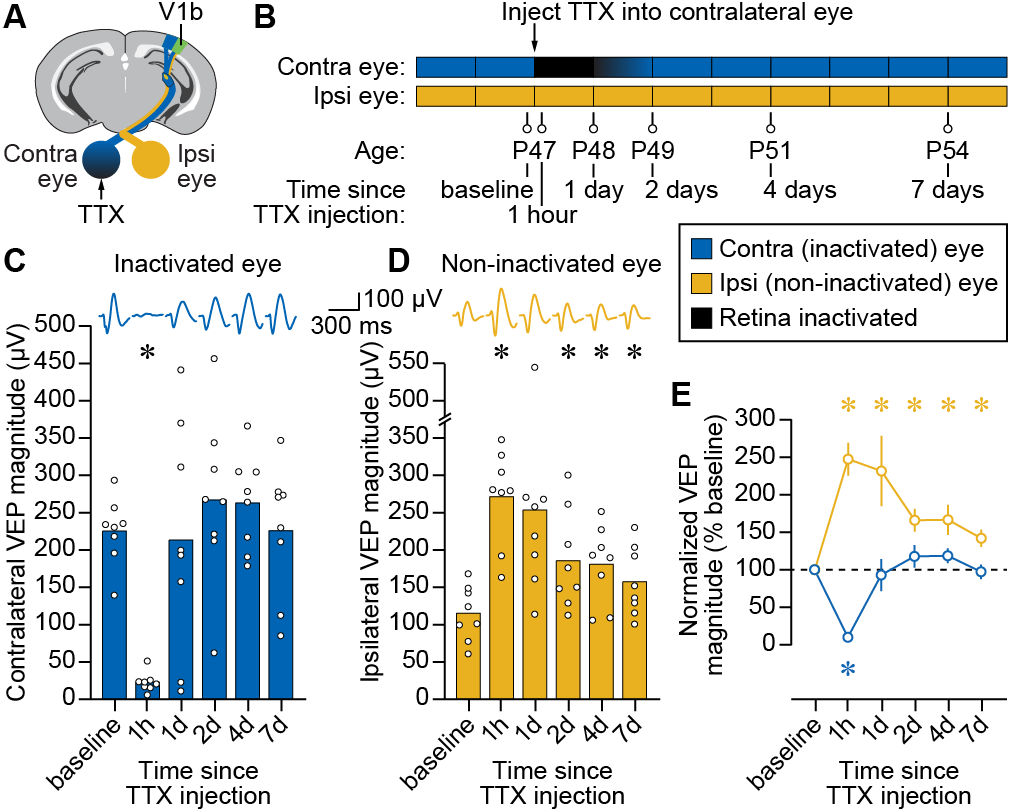
Temporary inactivation of one retina potentiates visual responses to stimulation of the non-inactivated eye in awake, late-adolescent mice. **(A)** Schematic of mouse brain showing recording site in V1b and TTX injection site into contralateral eye. **(B)** Experimental timeline showing ocular manipulation and VEP recording session times. **(C)** Longitudinal measurements of VEP magnitude for stimulation of the contralateral, inactivated eye. Filled bars denote mean peak-to-peak magnitude and open circles denote individual biological replicates. Average VEP waveforms are shown above for each time point. Asterisk denotes a significant difference in VEP magnitude (P<0.05) compared to the baseline recording. Analyses performed using one-way repeated measures ANOVA(F= 12.95, P=0.0013) followed by Dunnett’s multiple comparisons tests (compared to baseline: 1 h, P<0.0001; 1d, P=0.9987; 2d, P=0.6037; 2d, P=0.2974; 7d, P>0.9999). **(D)** Same as C for non-inactivated, ipsilateral eye. Analyses performed using one-way repeated measures ANOVA(F=9.361, P=0.0092) followed by Dunnett’s multiple comparisons tests (compared to baseline: 1 h, P=0.0001; 1d, P=0.0639; 2d, P=0.0111; 2d, P=0.0216; 7d, P=0.0148). **(E)** VEP magnitude overtime normalized to baseline. Blue trace denotes inactivated contralateral eye; yellow trace denotes non-inactivated ipsilateral eye; open circles denote mean; error bars denote SEM; blue and yellow asterisks denote significant differences (P<0.05) compared to hypothesized value of 100% for the contralateral and ipsilateral eyes, respectively. Analyses were performed using a one-sample t-test for the contralateral eye (compared to baseline: 1 h, P<0.0001; 1 d, P=0.7475; 2d, P=0.2739; 2d, P=0.0976; 7d, P=0.7856) and a one-sample Wilcoxon test for the ipsilateral eye (compared to baseline: 1 h, P=0.0078; 1d, P=0. 0078; 2d, P=0.0156; 2d, P=0. 0156; 7d, P=0. 0156), with test identity selected based on outcome of Shapiro-Wilk normality test. All data in this figure is for phase-reversing sinusoidal grating stimulation at a spatial frequency of 0.2 cpd.

### Fellow eye inactivation promotes stable recovery following long-term monocular deprivation in mouse V1

We next asked how temporary inactivation of the fellow eye would impact vision in amblyopic animals. To model deprivation amblyopia, we subjected mice to long-term MD initiated during the peak of the classical critical period for juvenile ocular dominance plasticity and extending two weeks beyond its closure (P26-47). To assess the consequences of fellow eye inactivation in amblyopic mice, we monitored VEPs from V1 just after opening the monocularly deprived eye, and then for several weeks after delivering a single intravitreal injection of TTX into the non-deprived eye (MD then TTX group; **Fig. 2A**). Littermate controls were distributed into two additional groups: one undergoing 3 weeks of MD followed by fellow eye saline injection (MD group), and another undergoing sham eyelid suture/re-opening followed by a fellow eye saline injection (Sham group). In all cases, the deprived (or sham deprived) eye was contralateral to the recording electrode while the injected non-deprived fellow eye was ipsilateral. Immediately following eye opening, both groups that underwent MD showed clear depression of deprived contralateral eye VEPs, with response magnitudes approximately half of those observed in sham controls (**Figs. 2B-C**). Longitudinal tracking of these animals revealed little change in contralateral visual responsiveness over several weeks in sham or MD animals receiving ipsilateral eye saline injections, confirming stability of the visual deficit. Strikingly, however, animals receiving fellow eye TTX injections following MD showed a significant increase in deprived eye responses (**Fig. 2B**) to values that were comparable to sham controls (**Fig. 2C**). The recovery of contralateral eye responses to normal levels was stable for many weeks (**Figs. 2B-C**) and occurred across a range of spatial frequencies (**Fig. 2D, Table S1A**). These results indicate that temporary inactivation of the fellow eye promotes rapid, complete and apparently permanent recovery of amblyopic eye responses in V1.

**Figure 2:**
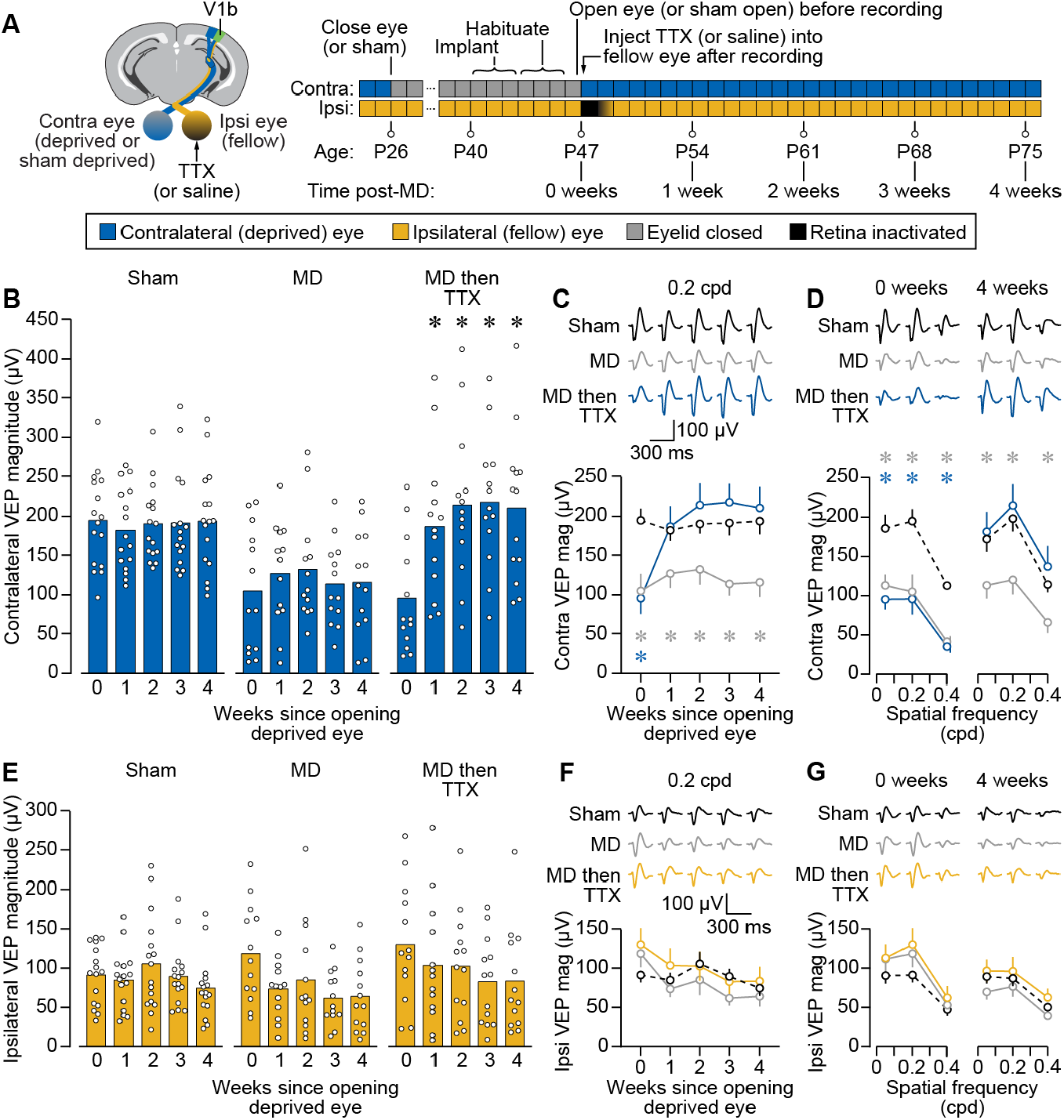
Fellow eye inactivation in mice promotes stable and complete recovery of vision in both eyes following long-term monocular deprivation. **(A)** Left, schematic of mouse brain showing recording site in V1b, as well indicating deprivation and inactivation in the contralateral and ipsilateral eyes, respectively. Right, timeline showing experimental manipulations and recording session times. **(B)** Longitudinal measurements of VEP magnitude for stimulation of the contralateral eye at 0.2 cpd for three littermate treatment groups: Sham, sham contralateral eyelid suture P26-47 and fellow eye saline at P47; MD, contralateral eyelid suture P26-47 and fellow eye saline at P47; MD then TTX, contralateral eyelid suture P26-47 and fellow eye TTX at P47. Filled bars denote mean peak-to-peak magnitude and open circles denote individual biological replicates. Asterisk denotes a significant difference in VEP magnitude (P<0.05) compared to before treatment (0 Weeks). Analyses were performed using a two-way repeated measures ANOVA (Treatment × Time, F(8,156)=6.925, P<0.0001) followed by Dunnett’s multiple comparisons tests (Sham, 0 vs. 1, 2, 3, 4 weeks: P=0.4963, 0.9774, 0.9986, 0.9999; MD, 0 vs. 1,2, 3, 4 weeks: P=0.5051,0.5160, 0.9756, 0.9640; MD then TTX, 0 vs. 1, 2, 3, 4 weeks: P=0.0052, 0.0024, 0.0007, 0.0010). **(C)** Contralateral VEP magnitude over time for stimulation at 0.2 cpd, with Sham in black (dashed), MD in grey, and MD then TTX in blue. Open circles denote mean, error bars denote SEM; average waveforms shown above plot; grey and blue asterisks denote significant difference compared to Sham (P<0.05) for the MD and MD then TTX groups, respectively, computed using Dunnett’s multiple comparisons tests (MD vs. Sham at 1, 2, 3, 4, 5 weeks: P=0.0050, 0.0414, 0.0356, 0.0025, 0.0077; MD then TTX vs. Sham at 1, 2, 3, 4, 5 weeks: P=0.0011, 0.9806, 0.6524, 0.5640, 0.8190). **(D)** Contralateral VEP magnitude across different spatial frequencies just after opening deprived eye but before treatment (0 weeks) and 4 weeks after opening deprived eye and inactivating fellow eye. Symbols/colors same as C. Post-hoc comparisons performed using Dunnett’s multiple comparisons tests (0 weeks, Sham vs. MD at 0.05, 0.2, 0.4 cpd: P=0.0054, 0.0050, 0.0002; 0 weeks, Sham vs. MD then TTX at 0.05, 0.2, 0.4 cpd: P=0.0006, 0.0011, <0.0001; 4 weeks, Sham vs. MD at 0.05, 0.2, 0.4 cpd: P=0.0311, 0.0077, 0.0157; 4 weeks, Sham vs. MD then TTX at 0.05, 0.2, 0.4 cpd: P=0.9319, 0.8190, 0.6413). **(E-G)** Same as B-D but for ipsilateral (fellow, inactivated) eye, with yellow denoting the MD then TTX condition in F-G. Analyses were performed using a two-way repeated measures ANOVA (Treatment × Time, F(8,156)=1.464, P=0.1745), with the absence of a significant interaction suggesting that there were no differences over time that could be attributed to treatment.

We also measured V1 responses to stimulation of the non-deprived fellow eye (**Table S1B**). Immediately after opening the deprived eye, response magnitudes through the non-deprived fellow eye were slightly elevated above those of sham animals (**Figs. 2E-G**), consistent with potentiation of non-deprived eye responses that has been well documented in rodent V1 (Frenkel & Bear, 2004). The initial potentiation of fellow eye responses returned to sham levels during the weeks after re-opening the deprived eye (**Figs. 2E-G**). These results show that unlike the stable response depression observed for the deprived eye following MD alone (**Figs. 2B-D**), potentiation of fellow eye responses following MD appears transient. In addition, the trajectory of fellow eye responses over time was not affected by the temporary retinal inactivation of this eye; responses recovered fully and were indistinguishable from sham controls during the weeks after re-opening the deprived eye irrespective of treatment (**Figs. 2E-G**). Together, these results demonstrate that following amblyogenic rearing in mice, fellow eye inactivation initiated beyond the juvenile sensitive period fosters a return of normal vision through both eyes.

### Reverse occlusion promotes transient recovery following long-term monocular deprivation in mouse V1

The standard treatment for amblyopia in infants and young children is to temporarily occlude vision through the fellow eye to promote vision through the amblyopic one. In animal models, this can be simulated using reverse occlusion (RO), wherein the fellow eye is temporarily sutured closed after the period of monocular deprivation. Reverse occlusion degrades visual responses to patterned stimuli but does not eliminate ganglion cell activity.

To directly compare the impact of fellow eye inactivation and reverse occlusion on the amblyopic visual cortex, we subjected littermate mice to long-term MD and then either briefly inactivated or sutured closed the fellow eye for 1 week (**Fig. 3A**). We controlled for the effect of injection and eyelid suture using saline injections in the RO group and sham eyelid closure/opening in the TTX group. These experiments replicated the finding that a single injection of TTX into the fellow eye promoted recovery of deprived eye responses that was stable for many weeks across a range of spatial frequencies (**Figs. 3B-D, Table S2A**). Interestingly, 1 week of RO also led to potentiated responses through the originally deprived eye. However, unlike fellow eye inactivation, the gains observed after RO were short-lived and dissipated over time, presumably because of the age at which it was initiated (P47). Responses to fellow eye stimulation were again elevated immediately after opening the deprived eye, and lessened in the weeks that followed (**Figs. 3E-G, Table S2B**). There was no significant difference in fellow eye response magnitude between treatment groups. Collectively, these results indicate that the plasticity driven by RO in adolescent mice is temporary, whereas the recovery driven by fellow eye inactivation is long lasting.

**Figure 3:**
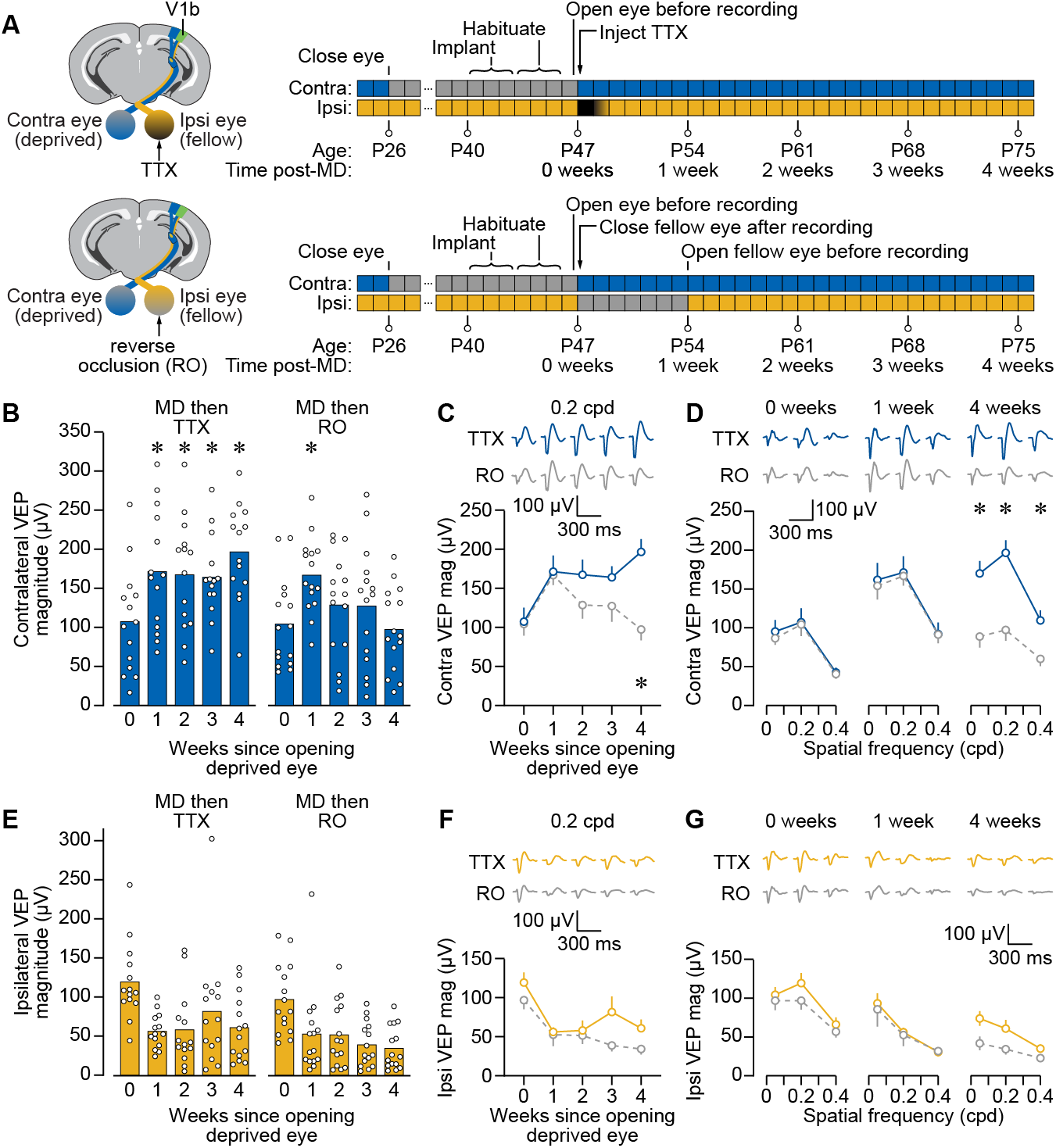
Reverse occlusion improves deprived eye vision following long-term monocular deprivation, but the recovery is not lasting. **(A)** Schematic and timelines for experiment comparing fellow eye inactivation to RO following 3 weeks MD. **(B-G)** Same format as Fig. 2B-G, but for two littermate treatment groups: MD then TTX, contralateral eyelid suture P26-47 and fellow eye TTX injection + sham suture at P47 + sham opening at P54; MD then RO, contralateral eyelid suture P26-47 and fellow eye saline + reverse occlusion at P47-54. MD then TTX group is blue for C-D and yellow for F-G. MD then RO group is grey (dashed) for C-D and F-G. Except where otherwise noted, all data in this figure are for phase-reversing sinusoidal grating stimulation at a spatial frequency of 0.2 cpd. Analyses in B and E were performed using two-way repeated measures ANOVA tests (Contralateral eye, Treatment × Time, F(4,108)=6.363, P=0.0001; Ipsilateral eye, Treatment × Time, F(4,108)=1.474, P=0.2152), with the significant interaction for the contralateral eye motivating Dunnett’s multiple comparisons tests (MD then TTX, 0 vs. 1, 2, 3,4 weeks: P=0.0057, 0.0065, 0.0437, 0.0042; MD then RO, 0 vs. 1, 2, 3, 4 weeks: P=0.0035, 0.4119, 0.5738, 0.9831). Dunnett’s post-hoc tests were also used for analyses in C (MD then TTX vs MD then RO at 0 and 1 weeks: P>0.9999, at 2, 3, 4 weeks: P=0.5361, 0.5446, 0.0004) and D (0 weeks, MD then TTX vs MD then RO at 0.05, 0.2, 0.4 cpd: P=0.9917, >0.9999, 0.9998; 1 week, MD then TTX vs MD then RO at 0.05, 0.2, 0.4 cpd: P=0.9995, >0.9999, >0.9999; 4 weeks, MD then TTX vs MD then RO at 0.05, 0.2, 0.4 cpd: P=0.0041, 0.0004, 0.0255).

### Fellow eye inactivation promotes stable recovery following long-term monocular deprivation in cat V1

We next evaluated fellow eye inactivation as a potential amblyopia treatment in the cat, the species that laid the foundation for much of what is known about ocular dominance plasticity and the pathogenesis of amblyopia. The electroencephalogram was recorded non-invasively over V1 during monocular viewing of phase-reversing grating stimuli at a range of spatial frequencies (**Fig. 4A**). The response to visual stimulation was quantified by calculating the total power at the 2 Hz phase reversal frequency and the next six harmonics (**Fig. 4B**). We compared these to the total power at control frequencies offset from the 2 Hz fundamental frequency and its harmonics by 0.45 Hz. As expected, the lowest spatial frequency presented (0.05 cpd) evoked the largest response. Using this approach, we were able to reliably detect responses to stimuli for spatial frequencies up to 0.5 cpd (**Fig. 4C**) with response profiles similar to time-domain VEP analysis (**Fig. S2**).

**Figure 4:**
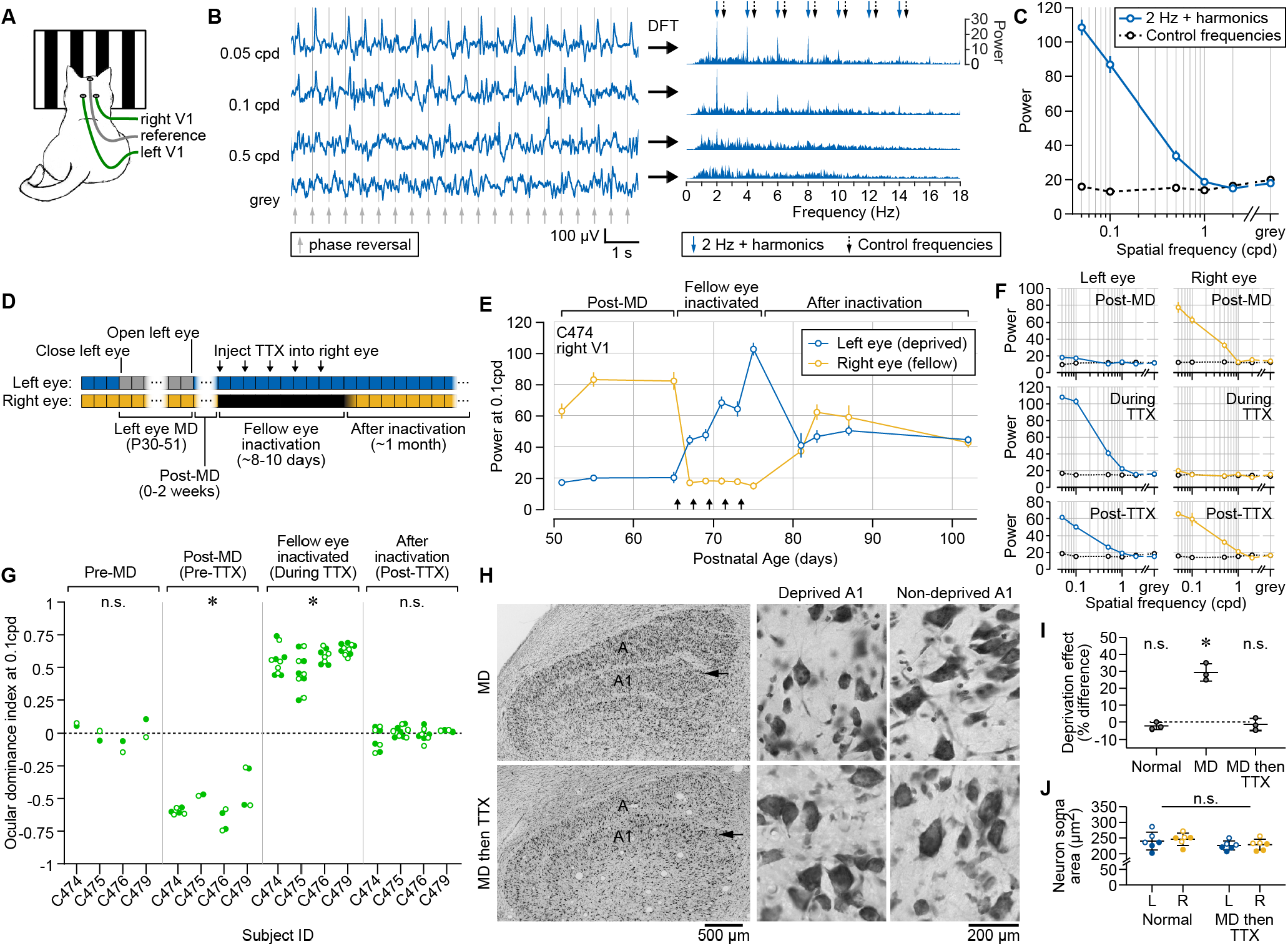
Fellow eye inactivation promotes stable functional and structural recovery in cats following long-term monocular deprivation. **(A)** Recording and visual stimulation setup for measuring visually-evoked responses non-invasively in cats. **(B)** Methodology for computing visually-evoked responses from scalp surface field potential. Left, example raw field potential time series recorded from V1 during presentation of phase-reversing visual stimuli at a range of spatial frequencies. Grey lines denote timing of phase reversals. Right, data on left shown in the frequency domain following discrete Fourier transform. Blue arrows point to peaks in spectral power at 2 Hz (the phase reversal frequency) and six harmonics, representing visually-evoked responses. Black arrows point to the frequencies used for control power measurements. **(C)** Total power at the visually-driven (phase reversing) frequency and its harmonics (blue) versus control (black) across a range of spatial frequencies. Values computed from same recording shown in B. **(D)** Timeline showing experimental manipulations and recording session times. **(E)** Total power of visually-evoked responses for one animal (C474, right V1) viewing visual stimuli through the deprived left eye (blue) versus the fellow right eye (yellow). Arrows denote time of TTX injections. Error bars, SEM. Spatial frequency, 0.1 cpd. **(F)** Total power of visually-driven responses for the same animal shown in E across a range of spatial frequencies before (top), during (middle), and after (bottom) inactivation of the fellow eye with TTX. Deprived left eye shown in blue, and fellow right eye shown in yellow. Black symbols denote control frequencies as in C. **(G)** Ocular dominance indices calculated for 4 cats before MD, after MD (but before inactivation), during inactivation, and after the fellow eye was no longer inactivated. Data are shown for both right (closed circles) and left (open circles) V1. Positive and negative values indicate response biases toward the left and right eyes, respectively, while the dashed line at 0 indicates balanced V1 responses expected from a neurotypical animal. As a reference, indices for the right hemisphere of C474 were calculated from data shown in E. Asterisks denote significant differences (p>0.05) from the hypothesized value of 0, analyzed using Wilcoxon signed-rank tests (Pre-MD, P=0.9453; Post-MD, P<0.0001; Fellow eye inactivated, P<0.0001; Post-TTX, P=0.8236). **(H)** Left, low magnification image of the LGN stained for Nissl substance after 3 weeks MD (top), or 3 weeks MD followed by fellow eye inactivation (bottom). Arrow indicates lamina A1, ipsilateral to the deprived eye. Middle and right, high magnification images from deprived (middle) and non-deprived (right) A1 layers for the same conditions shown on left. **(I)** Deprivation effect, stereological quantification of neuron soma size within deprived and non-deprived A and A1 layers, for normally-reared cats, cats undergoing 3 weeks of MD, and cats undergoing 3 weeks of MD followed by fellow eye inactivation. Asterisk denotes significant difference (p>0.05) from the hypothesized value of 0%, analyzed using a one-sample t-test (Normal, P=0.1917; MD, P=0.0100; MD then TTX, P=0.5716). **(J)** Average soma size for neurons in LGN layers downstream of the deprived left eye (L) or the fellow right eye (R) for normally-reared animals compared to those undergoing 3 weeks of MD followed by fellow eye inactivation. There was no significant difference between the groups (Welch’s ANOVA, W=1.426, P=0.2881). Open and closed symbols indicate values drawn from laminas A1 and A, respectively. Black lines indicate mean and SEM for each group.

We tracked longitudinally the visually-evoked responses in cats that had undergone 3 weeks of MD until P51, an age at which RO is minimally effective at reversing the effects of MD (Blakemore & Van Sluyters, 1974). After confirming the stable loss of visual responsiveness to the deprived eye, fellow eye inactivation was achieved over 8-10 days using intravitreal TTX (**Figs. 4D-F**). As expected, before MD all cats had balanced responses to stimulation of either eye, and this was strongly shifted after MD (**Figs. 4G, S3**) due to weaker responses to deprived eye stimulation (**Figs. 4E-F**). In three of the four animals, we allowed 1-2 weeks of binocular vision prior to initiating inactivation and verified that the impairment in deprived eye responses was stable. During the period of inactivation, fellow eye responses were reduced to the level of control frequencies and grey screen values, as expected (**Figs. 4F**). Meanwhile, responses to stimulation of the previously deprived (non-inactivated) eye were gradually potentiated (**Fig. 4E-F**). After the TTX wore off, responses to stimulation of the fellow eye returned, and we again observed balanced V1 responses to stimulation of both eyes that persisted for many weeks without sign of regression (**Figs. 4E-G, S3**). These observations were consistent across all four animals (**Fig. 4G**) at spatial frequencies up to the 0.5 cpd detection limit (**Table S3**), and there was no discernable decrement in efficacy for the animal that underwent fellow eye inactivation at the oldest postnatal age (P65; **Figs. 4E-F, S3A; Table S3A**). Post-mortem analysis confirmed normal ocular histology in all animals, consistent with previous observations (DiCostanzo, Crowder, Kamermans, & Duffy, 2020). These results demonstrate that, following amblyogenic rearing in cats, fellow eye inactivation promotes a balanced recovery in V1 downstream of both eyes, and this intervention is efficacious at ages beyond previously established sensitive periods for reversal. Although quantitative behavioral assessment was not performed, qualitative observation of the kittens showed unambiguously that vision and visually guided behavior was restored by treatment.

### Fellow eye inactivation corrects anatomical effects of monocular deprivation in cat dLGN

In cats and primates, inputs from the two eyes are distributed to different laminae of the dorsal lateral geniculate nucleus (dLGN). In these highly visual species, an anatomical hallmark of monocular deprivation is the reduction of dLGN neuron size within laminae downstream of the deprived eye (Duffy & Slusar, 2009; Wiesel & Hubel, 1963), a phenomenon associated with shrinkage of ocular dominance columns (Hubel, Wiesel, & LeVay, 1977) and dependent on synaptic modification in V1 (Bear & Colman, 1990). To explore the effect of fellow eye inactivation on an anatomical marker of ocular dominance plasticity, we compared post-mortem analysis of dLGN soma size from three cats that had undergone 3 weeks of MD followed by fellow eye inactivation with age-matched controls. Animals that underwent MD alone showed the classic shrinkage of dLGN cells in deprived laminae compared with the non-deprived lamina in the binocular segment (**Fig. 4H-I**). In contrast, cats subjected to MD that additionally underwent fellow eye inactivation showed comparable soma sizes in deprived and non-deprived dLGN lamina (**Figs. 4H-I**). Importantly, this was not the result of atrophy of cells downstream of both eyes, as the soma size of animals undergoing fellow eye inactivation was statistically indistinguishable from those of normally reared controls (**Fig. 4J**). These results provide evidence that, in addition to physiological recovery from MD (**Figs. 2B-G, 3B-G, 4E-G, S3; Table S3**), fellow eye inactivation reverses an anatomical correlate of ocular dominance plasticity in the cat visual pathway.

### Discussion

In animals, long-term monocular deprivation models the cause of the most severe form of human amblyopia, for which treatment options are absent or limited. Traditional therapies, such as patching the fellow eye, are beneficial only when initiated at very young ages (Birch & Stager, 1996). Even when patching is initially successful, recurrence of amblyopia is common (Bhola, Keech, Kutschke, Pfeifer, & Scott, 2006; Holmes et al., 2004). These observations supported the view that the brain loses plasticity shortly after birth, when synaptic connections are essentially set for life (LeVay, Wiesel, & Hubel, 1980). However, experiments over the past 40 years in a number of species have shown repeatedly that under certain conditions synaptic connections in V1 remain mutable across the lifespan. Notwithstanding the enormous potential these findings have for the treatment of amblyopia, efforts to translate this knowledge to human benefit have so far been unsuccessful. The steep challenge that remains is to devise a strategy for harnessing this plasticity to promote recovery of function that can be applied in a clinical setting.

The current study was motivated by numerous human clinical reports that stable and severe amblyopia can sometimes remit in adults when the fellow eye is damaged or removed (El Mallah et al., 2000; Kaarniranta & Kontkanen, 2003; Klaeger-Manzanell et al., 1994; Rahi et al., 2002; Vereecken & Brabant, 1984). Our experiments were designed to test the hypothesis that this remission is enabled, not by permanent removal of interocular suppression by the fellow eye, but by eliminating activity in the eye only for as long as required to enable connections from the amblyopic eye to become reestablished in V1. The data, obtained in two evolutionarily distant animal models, clearly indicate that stable and severe amblyopia can be reversed following temporary inactivation of the fellow eye. This recovery from amblyopia is durable, persisting for weeks after the TTX has worn off and vision is fully restored in the inactivated eye, and is observed in animals at ages beyond the classically defined critical period.

The current findings can be understood in the context of “Hebbian” models of synaptic strengthening. By inactivating the dominant input, homeostatic adaptations occur in V1 that enable visual experience to drive potentiation of synapses that are otherwise too weak to be modified (see, e.g., (Clothiaux, Bear, & Cooper, 1991)). In this conceptual framework, recovery might be further enhanced as activity in the fellow eye returns, analogous to the phenomenon of associative long-term potentiation (Barrionuevo & Brown, 1983). We note that this additional bootstrapping of the weak eye input would not be available if the fellow eye were permanently removed, perhaps explaining why in humans loss of the fellow eye leads to recovery from amblyopia in only ∼10% of cases after age 11 (Rahi et al., 2002). We are far more successful in reversing amblyopia after the critical period in the animal models using temporary inactivation of the fellow eye than what has been observed using enucleation (Drager, 1978; Harwerth et al., 1984; Kratz & Lehmkuhle, 1983).

The end of the critical period is typically defined as the age when RO is no longer able to reverse the effects of early MD. In mice, the critical period has been estimated to end at approximately P32 (Gordon & Stryker, 1996), so we were surprised to see that seven days of RO initiated at P47 restored responses through an eye that had been rendered amblyopic with 3 weeks of MD. However, unlike what was observed with fellow eye inactivation, the improvement after RO was short lived. Similar reemergence of visual impairment following RO has been reported in amblyopic kittens (D. E. Mitchell, Murphy, & Kaye, 1984). It is noteworthy that although occlusion therapy is the standard of care for visual correction in children, approximately one-quarter of patients experience worsening vision through the amblyopic eye within a year after treatment concludes (Bhola et al., 2006; Holmes et al., 2004). These clinical findings highlight the importance of evaluating the long-term stability of recovery in preclinical studies. While a number of potential therapies for amblyopia have been identified in rodent models, long-term stability is rarely monitored.

Occlusion and retinal inactivation are qualitatively different manipulations of visual experience, and have profoundly different consequences on the visual cortex. Eye patches and eyelid sutures degrade image form, but have little effect on the overall activity of retinal ganglion cells (Kuffler, Fitzhugh, & Barlow, 1957). Instead, occlusion replaces spatiotemporally structured patterns of activity with stochastic noise. During early development, it has been shown that this retinal noise can rapidly trigger the synaptic depression in V1 that causes amblyopia (Cooke & Bear, 2014). In contrast, visual responses and thalamocortical synaptic strength are preserved in V1 following a comparable period of intravitreal TTX (Coleman et al., 2010; Frenkel & Bear, 2004; Iurilli, Benfenati, & Medini, 2012; Rittenhouse, Shouval, Paradiso, & Bear, 1999). Although RO can promote recovery during the critical period, our experiments show that only inactivation produces a durable effect when initiated later in life.

Previous studies in cats and rodents showed that recovery from a brief period of monocular deprivation could be promoted by prolonged immersion in complete darkness as well as by temporary bilateral retinal inactivation (Duffy & Mitchell, 2013; Fong, Mitchell, Duffy, & Bear, 2016; He, Ray, Dennis, & Quinlan, 2007). It is possible that dark exposure, binocular TTX, and fellow eye inactivation all tap into a common mechanism, but there are some important differences. To be effective, the period of darkness must be long (≥10 days) and cannot be interrupted, even briefly, by light exposure (D. E. Mitchell, MacNeill, Crowder, Holman, & Duffy, 2016). Treatment of both retinas with TTX overcomes some of these limitations, but so far has been shown only to reverse the effects of short-term MD (Fong et al., 2016). Darkness and binocular TTX are also less effective than fellow eye inactivation in promoting anatomical recovery beyond the critical period (Duffy, Fong, Mitchell, & Bear, 2018). Furthermore, from a clinical standpoint, total visual deprivation is not practical. These procedures are known to disrupt circadian rhythms and cause visual hallucinations (L. Pang, 2016), and would necessitate continuous patient care and supervision. Thus, finding that monocular TTX treatment is effective in reversing amblyopia caused by long-term MD in both cats and mice represents a substantial advance.

The precise mechanisms that enable connections downstream of the amblyopic eye to gain strength following fellow eye inactivation remain to be determined. However, examining the published consequences of monocular inactivation and related manipulations may provide some insight. Similar to our observation that retinal inactivation significantly potentiates responses through the non-inactivated eye, both monocular enucleation and optic nerve crush in adult rodents augment the spared eye responses (Nys et al., 2014; Vasalauskaite, Morgan, & Sengpiel, 2019). Unilateral retinal inactivation and enucleation both reduce markers of synaptic inhibition in V1 of monkeys (Hendry, Fuchs, deBlas, & Jones, 1990; Hendry et al., 1994; Hendry & Jones, 1988) and mice (Barnes et al., 2015; Maffei & Turrigiano, 2008). A change in inhibition, even if transient, could be sufficient to promote Hebbian potentiation of excitatory synapses serving the amblyopic eye. Excitatory synaptic plasticity could also be facilitated by observed increases in principal cell excitability (Barnes et al., 2015; Maffei & Turrigiano, 2008) and adjustments in NMDA receptor properties (Philpot, Espinosa, & Bear, 2003; E. M. Quinlan, Olstein, & Bear, 1999) that have been observed after dark exposure, retinal inactivation, or enucleation. Any or all of these homeostatic adjustments after fellow-eye inactivation could account for the observed recovery of vision after the critical period, and it will be of interest in future studies to pinpoint the essential modifications. Although we do not yet know the precise mechanism, this gap in knowledge does not preclude application of insights gained here to amblyopia treatment. The mechanism for recovery enabled by patch therapy during the critical period is still not known, and yet this is the current standard of care for amblyopia.

The strengths of the current study are (1) that it is one of the few to satisfy the “two-species” rule for establishing potential clinical efficacy, (2) it distinguishes between two competing hypotheses for how adult enucleation of the fellow eye enables recovery from amblyopia, and (3) the experiments demonstrate durable recovery from the effect of long-term MD after the end of the critical period. An injection of TTX (or equivalent) into the eye is not without risks that need to be carefully considered before contemplating human application (see, e.g., (Dossarps et al., 2015)). However, histological analysis of the cat retina and optic nerve revealed no deleterious effect of 10 days of TTX treatment (DiCostanzo et al., 2020), and the electrophysiological analyses and behavioral observations in the current study indicate complete functional recovery of vision in the injected eye. Similarly, experiments in awake, behaving monkeys show full recovery of visual acuity and eye reflexes following intravitreal TTX (Foeller & Tychsen, 2019). Complete retinal inactivation in cats and monkeys can be achieved with 1.5-15 µg of TTX confined to the vitreous humor (Ataman et al., 2016; DiCostanzo et al., 2020). In humans, 30 µg of TTX administered subcutaneously twice a day for 4 days was reported to be safe and well tolerated (Hagen et al., 2017). In addition, new biodegradable polymers have recently been developed for therapeutic delivery of TTX to achieve prolonged and local sodium channel blockade without detectable systemic toxicity (Zhao et al., 2019). Thus, there may be a path forward to apply this strategy for the treatment of deprivation amblyopia in adult patients.

## Materials and Methods

### Mouse studies

Mouse experiments were conducted using male and female wildtype animals on the C57BL6/N background. Animals were purchased from Charles River for experimentation or as breeders for a colony maintained at MIT. Offspring were housed in groups of 2-5 same-sex littermates after weaning at P21 and maintained on a 12h light-12h dark cycle. All recordings were conducted during the light cycle. Food and water were available *ad libitum*. Rearing and experimental procedures were conducted in accordance with protocols approved by the Committee on Animal Care at MIT, and conformed to guidelines from the National Institutes of Health and the Association for Assessment and Accreditation of Laboratory Animal Care International.

#### Experimental Design

For all experiments, chronic electrode implant surgery was conducted between P40 and P44, and animals were habituated to head fixation and monocular grey screen viewing on two separate days between P42 and P46. For monocular inactivation experiments (**Fig. 1**), baseline recordings were conducted followed by intravitreal TTX injections into the eye contralateral to the recording electrode. Recordings were then conducted 1 hour, 24 hours, 48 hours, 4 days, and 7 days after the injection. Target sample size (number of biological replicates) was computed in SPSS for a power of 0.85 and effect size measured in pilot data (not included in study). For MD experiments (**Figs. 2-3**), eyelid suture (or sham suture) was performed at P26. Deprived eyes were re-opened (or sham re-opened) at P47 (5-7 days after implant surgery), and the first electrophysiological recording from V1 took place 45 minutes later. Shortly after the first recording session, animals underwent either intravitreal injections of TTX or saline into the fellow eye, in some experiments with RO or sham RO. Recording sessions occurred weekly thereafter until P75. All recordings were conducted blind to deprivation and treatment conditions, and each littermate group was handled by the same experimenter each week. Target sample size was estimated based on previously published data (Fong et al., 2016). At the conclusion of the experiment, mice were euthanized, and the brains and eyeballs were harvested for post-mortem analyses. In order to be included in the final data set, mice needed to meet all of the following *a priori* determined inclusion criteria: mice maintained good health status throughout experiment; eyelid stayed fully closed during MD and RO; a visually-evoked response greater than noise was detectable through at least 1 eye at baseline; cornea, retina, and all parts of eyeball were healthy and intact; an electrode track was observable within L4 of binocular V1; at least 1 other same-sex littermate was included in another experimental group. Sample sizes reported refer to biological replicates (distinct animals used to capture the biological variation associated with each condition). Individual animals were tracked longitudinally, although technical replicates (independent repeated measurements at each time point) were not used.

#### Eyelid suture

Animals were anesthetized via inhalation of isoflurane (1-3% in oxygen), and body temperature was maintained on a heated surface at 37° C. Both eyes were moistened with sterile saline. For monocular deprivation and reverse occlusion, the eye being sutured first had fur on upper and lower eyelid of trimmed, and the eyeball surface rinsed with sterile saline. A thin layer of ophthalmic ointment containing bacitracin, neomycin, and polymysin was placed on the eye. The eyelid was closed using a single mattress stitch of polypropylene suture (7-0 Prolene), with care taken not to touch the corneal surface, and a square knot was tied on the exterior of the lower lid. Ophthalmic ointment was applied where the suture needle passed through the eyelid. Nails on the forepaws were filed or lightly trimmed. For sham sutures, the suture was cut and removed just prior to turning off anesthesia. To remove sutures (3 weeks later for MD and 1 week later for RO), mice were anesthetized and maintained as previously described. The suture was cut and removed, and eyelids were gently separated. The corneal surface was rinsed with sterile saline and examined for signs of damage. Sham animals were anesthetized, placed under the bright microscope for an equivalent amount of time as MD and RO animals, and had their sham MD or RO eyeball rinsed. Animals recovered from MD, RO, and eye opening procedures in their home cages.

#### Intravitreal injections

Mice were anesthetized using isoflurane, maintained at 37° C, and had both eyes moistened thereafter as described for eyelid sutures. For the injected eye, a ∼500 nm incision was made at the temporal corner of the eye, and a sterile 7-0 silk suture thread was pulled through the exposed sclera. The suture was pulled toward the nasal aspect to secure the globe and expose the temporal aspect of the eyeball. A 30-gauge needle was used to penetrate to the sclera and globe. The eye was rinsed with sterile saline just prior to inserting a glass micropipette into the vitreous chamber. A nanoliter injector was used to deliver 1 µl of either TTX (1 mM in citrate buffer) or saline. The micropipette was removed 1 minute after the injection. The eye was rinsed with sterile saline and the spot of penetration was coated with ophthalmic ointment containing bacitracin, neomycin, and polymysin. Animals recovered from intravitreal injection in their home cages.

#### V1 implant surgery

Pre-operative buprenorphine (0.1 mg/kg s.c.) was administered, and mice were subsequently anesthetized (1-3% isoflurane) and maintained at 37° C. Ophthalmic ointment was applied to both eyes (or on the eyelid surface for eyes that were sutured closed). The fur on top of the head was shaved and the exposed scalp was cleaned using ethanol (70% v/v) and a povidone-iodine solution (10% w/v). An incision was made the midline and connective tissue on the skull surface was removed. A steel post was affixed to the skull anterior to Bregma. A craniotomy was made over the right prefrontal cortex for implanting a silver wire reference electrode on the cortical surface. Another craniotomy was made over binocular V1 (3 mm lateral of Lambda) contralateral to the monocularly deprived (or sham MD) eye, and a tungsten microelectrode (300-500 kΩ; FHC 30070) was lowered to L4 (450 µm from cortical surface). The steel posts and male gold pins coupled to electrodes were secured to the skull using cyanoacrylate. The skull was then covered in dental cement (Orthojet). Animals were removed from anesthesia and transferred to a heated recovery chamber. Meloxicam (1 mg/kg s.c.) was administered during this initial recovery phase, as well as for 2 days post-operatively. Health was carefully monitored by experimenters and veterinary staff, and supplemental fluids or heat was delivered if needed.

#### Electrophysiological recording

All mouse recordings were conducted in awake, head-fixed animals with full-field visual stimuli presented on an LCD monitor in the binocular visual field at a viewing distance of 20 cm. On the days prior to the initial recording, animals were habituated to head restraint, grey screen viewing, and an opaque occluder that restricted vision to one eye or the other. On each recording day, local field potential (LFP) data were recorded from layer 4 of binocular V1 during visual stimulation sessions lasting 13.5 minutes per eye. Continuous LFP data were collected, amplified, digitized and low-pass filtered using the Recorder-64 system (Plexon). Other than the visual stimulus, the experimental room was kept dark and free of distractions (e.g. neither the experimenter nor other mice were present in the room during the recording session). Stimuli were generated using custom software written in MATLAB using the PsychToolbox extension (Brainard, 1997; Pelli, 1997), and consisted of sinusoidal oriented gratings phase reversing at 2 Hz. Stimuli were presented in blocks consisting of 50 phase reversals at 100% contrast for each of the following pseudo-randomly presented spatial frequencies: 0.05, 0.2, 0.4 cycles per degree (cpd). Each block of stimuli was separated by a 30-second presentation of a luminance-matched grey screen. A distinct orientation (offset by at 30° offset from any previously viewed orientation) was used each week, and only non-cardinal orientations were selected for presentation. All recordings were performed blind to treatment condition.

#### Electrophysiological data analysis

The VEP was defined as the phase reversal-triggered LFP, averaged across all phase reversals within a recording session for each individual animal, time point, and viewing eye. VEP waveforms and amplitudes were extracted from recorded LFP data using software developed by Jeffrey Gavornik (github.com/jeffgavornik/VEPAnalysisSuite). VEP magnitude was defined as the peak-to-peak amplitude of each biphasic VEP waveform. Data for the intermediate spatial frequency (0.2 cpd) is used for visualization, but data for lower and higher spatial frequencies is provided Tables S1-2.

#### Post-mortem analyses

Mice were deeply anesthetized using isoflurane and rapidly decapitated. Both eyeballs were removed for immediate ocular examination and dissection in phosphate-buffered saline (PBS). The corneas were examined under a microscope for signs of damage, and parts of the eye were thereafter dissected to look for abnormalities, particularly of the lens or retina. Meanwhile, the brains were harvested and fixed in 4% paraformaldehyde at room temperature. Tissue remained in fixation solution for 72 hours, and was thereafter stored in PBS. Coronal slices of the V1 were made at 50 µm using a vibrating microtome. Slices were rinsed with water and phosphate buffer, and then mounted on charged glass slides. The mounted slices dried at room temperature and ambient humidity for 24 hours. Nissl bodies were then stained using cresyl violet, and coverslipped with 1.5 glass and touline-based mounting medium (Permount). Slices were imaged on a confocal microscope (Olympus) using the transmitted light channel, and digital micrographs of electrode tracks were saved. Micrographs were compared to a mouse brain atlas to determine localization of electrode tracks in layer 4 of binocular V1. All analyses described were performed blind to treatment condition.

### Cat studies

Physiological and anatomical studies were conducted on 9 male and female cats that were all born and raised in a closed breeding colony at Dalhousie University. Rearing and experimental procedures were conducted in accordance with protocols approved by the University Committee on Laboratory Animals at Dalhousie University, and conformed to guidelines from the Canadian Council on Animal Care.

#### Experimental Design

Animals were monocularly deprived for three weeks starting at the peak of the critical period (P30). Four animals subjected to physiological assessment had their deprived eye opened and were administered 4-5 intraocular injections of TTX into the fellow eye immediately (C475), or following 1 (C476, C479) or 2 weeks (C474) of binocular vision. Animals underwent electroencephalogram recordings at various points throughout the experimental timeline. The number of animals used in the study was determined in consultation with veterinary staff, with 3 littermates being tested first in parallel, and 1 animal being tested later to validate results in an age-matched animal from a different litter. Three animals (C474, C475, C476) were euthanized at ∼P110, and tissue was harvested for anatomical analysis. Anatomical assessments were made on two additional groups of animals, which acted as our controls. The first control group (n=3) was normally reared until age 14 weeks and then received 5 injections of vehicle solution (citrate buffer) into the right eye. The second control group (n=3) was monocularly deprived at P30 for three weeks (n=3). Sample sizes of control groups for anatomy were designed to match the number of animals in the treatment group.

##### Eyelid suture

Monocular deprivation was performed under general gaseous anesthesia (3-4% isoflurane in oxygen) and involved closure of the upper and lower palpebral conjunctivae of the left eye with sterile polyglactin 910 thread (Vicryl), followed by closure of the eyelids with silk suture. Following the procedure, animals were administered Metacam (0.05 mg / kg) for post-procedure analgesia, local anesthesia was produced with Alcaine sterile ophthalmic solution (1% proparacaine hydrochloride; CDMV, Canada), and a broad-spectrum topical antibiotic (1% Chloromycetin; CDMV) was administered to mitigate infection after surgery.

##### Intravitreal injections

Upon completion of the 3-week MD period, animals were anesthetized with 3-4% isoflurane and the eyelids were opened. Either immediately or after a period of binocular vision, animals had their fellow eye inactivated with intravitreal injection of TTX (ab120055; abcam, USA) that was solubilized in citrate buffer at 3mM. Dosage was 0.5 µl / 100 gram body weight but irrespective of weight, injection volume never exceeded 10 µl per injection. This approximate dosage blocks action potentials of affected cells without obstructing critical cellular functions such as fast axoplasmic transport (Ochs & Hollingsworth, 1971). Injections were administered through a puncture made with a disposable sterile needle that created a small hole in the sclera located at the pars plana. Using a surgical microscope, the measured volume of TTX solution was dispensed into the vitreous chamber with a sterilized Hamilton syringe (Hamilton Company, USA) fixed with a 30 gauge needle (point style 4) that was positioned through the original puncture and about 5-10 mm into the chamber angled away from the lens. The total volume of TTX was dispensed slowly, and when complete the needle was held in place for about a minute before it was retracted. Following intraocular injection, topical antibiotic (1% Chloromycetin) and anesthetic (Alcaine) were applied to the eye to prevent post-injection complications, and Metacam (0.05 mg / kg) was administered for post-procedure analgesia. Animals received 4-5 injections, one every 48 hours, and for each injection the original puncture site was used to avoid having to make another hole. During the period of inactivation, we employed basic assessments of visual behavior and measured VEPs to confirm inactivation. We verified the absence of a pupillary light reflex as well as the lack of visuomotor behaviors such as visual placing, visual startle, and the ability to track a moving laser spot. These assessments were made while vision in the non-injected eye was briefly occluded with an opaque contact lens.

##### Electrophysiological recording

All cat recordings were conducted in anesthetized animals with full-field visual stimuli presented on an LCD monitor in the binocular visual field at a viewing distance of 70 cm. In preparation for each recording session, animals were anesthetized with 1-1.5% isoflurane, and supplemental sedation was provided with intramuscular acepromazine (0.06-0.1mg/kg) and butorphanol (0.1-0.2 mg/kg). Hair on the head was trimmed and a disposable razor was used to shave parts of the scalp where recording sites were located, two positioned approximately 2-8 mm posterior and 1-4 mm lateral to interaural zero over the presumptive location of the left and right primary visual cortices, and another site over the midline of the frontal lobes that acted as a reference. Electrode sites were abraded with Nuprep EEG skin preparation gel (bio-medical, MI, USA), and were then cleaned with alcohol pads. Reusable 10 mm gold cup Grass electrodes (FS-E5GH-48; bio-medical) were secured to each electrode site using Ten20 EEG conductive paste (bio-medical, USA) that was applied to the scalp. Impedance of the recording electrodes was measured in relation to the reference electrode to ensure values for each were below 5 kΩ. Electrophysiological signals were amplified and digitized with an Intan headstage (RHD2132; 20kHz sampling frequency), then recorded using an Open Ephys acquisition board and GUI software (Open Ephys, USA)(Siegle et al., 2017). Stimuli were generated using custom software developed in Matlab by Nathan Crowder and Braden Kamermans using the Psychophysics Toolbox extension (Brainard, 1997; Pelli, 1997), and consisted of full contrast square wave gratings with a 2 Hz contrast reversal frequency (Bonds, 1984; Norcia, Appelbaum, Ales, Cottereau, & Rossion, 2015; X. D. Pang & Bonds, 1991). Blocks of grating stimuli at different spatial frequencies (0.05, 0.1, 0.5, 1, and 2 cpd) or a luminance-matched grey screen were presented in pseudo-random order for 20 s each, with the grey screen also displayed during a 2 s interstimulus interval. Each block was repeated at least 6 times. Each eye was tested in isolation by placing a black occluder in front of the other eye during recording. Eyes were kept open with small specula, and the eyes were frequently lubricated with hydrating drops. Recording sessions lasted about 1 hour and animal behavior was observed for at least an additional hour post-recording to ensure complete recovery.

##### Electrophysiological data analysis

The raw electroencephalogram was imported to MATLAB where it was high-pass filtered above 1 Hz, then subjected to Fourier analysis (Bach & Meigen, 1999; Norcia et al., 2015). The magnitude of VEPs was calculated as the sum of power at the stimulus fundamental frequency plus 6 additional harmonics (2-14 Hz) (DiCostanzo et al., 2020). Baseline nonvisual activity was calculated as the sum of power at frequencies 0.2Hz offset from the visual response (2.2-14.2Hz). Parallel analysis of EEG data was performed using the EEGLab and ERPLab toolboxes for MATLAB (Black et al., 2017; Delorme & Makeig, 2004; Lopez-Calderon & Luck, 2014). The data was bandpass filtered, segmented, and normalized for extracting event-related potentials for time-domain VEP analysis. Response profiles were similar using frequency- and time-domain analyses (**Figs. 4C, S2B**), but variability in peak VEP latencies and polarities after MD and during inactivation made frequency domain analysis a more practical choice for longitudinal comparisons. Ocular dominance index was computed as (*L-R*) / (*L+R*), where *L* is the mean summed power through the left (deprived) eye and *R* is the mean summed power through the right (fellow) eye. Data for the intermediate detectable spatial frequency (0.1 cpd) is used for visualization, but data for lower and higher spatial frequencies is provided **Table S3**.

##### Histology

In preparation for histology, animals were euthanized with a lethal dose of sodium pentobarbital (Pentobarbital Sodium; 150 mg/kg) and shortly thereafter exsanguinated by transcardial perfusion with approximately 150 ml of phosphate buffered saline (PBS) followed by an equal volume of PBS containing 4% dissolved paraformaldehyde. Brain tissue was immediately extracted and the thalamus was dissected from the remainder of the brain in order to prepare the LGN for sectioning and histological processing. Tissue containing the LGN was cryoprotected and then cut coronally into 25-µm thick sections using a sliding microtome. A subset of sections was mounted onto glass slides and stained with a 1% Nissl solution (ab246817; Abcam, USA), and then were coverslipped with mounting medium (Permount). To analyze cell size, the cross-sectional area of neuron somata within A and A1 layers of the left and right LGN was measured from Nissl-stained sections using the nucleator probe from a computerized stereology system (newCAST; VisioPharm, Denmark). All measurements were performed using a BX-51 compound microscope with a 60X oil-immersion objective (Olympus; Markham, Ottawa, Canada). Neurons were distinguished from glial cells using established selection criteria (Wiesel and Hubel, 1963) that included measurement of cells with dark cytoplastmic and nucleolar staining, and with light nuclear staining. Adherence to these criteria permitted avoidance of cell caps and inclusion only of neurons cut through the somal midline. Approximately 500-1000 neurons were measured from each animal. For each animal, an ocular dominance index was computed using the average soma size from each LGN layer: ((Left A1+Right A)-(Left A+Right A1) / (Left A1+Right A), which indicated the percentage difference between eye-specific layers.

#### Statistical Analyses

Statistical analyses were performed using Prism (GraphPad). Normality testing was performed using the Shapiro-Wilk, and the outcome was used to determine whether to proceed with parametric or nonparametric tests. Variance of all data sets was also computed and used to determine whether to select a test that assumed equal variances or not. Comparisons to noise levels (to meet inclusion criteria) were performed using a paired t-test (parametric) or Wilcoxon matched-pairs signed rank test (nonparametric). Comparisons of to a hypothetical mean (e.g. for normalized data) were performed using one-sample t-tests (equal variances) or Welch’s ANOVA (unequal variances). To analyze differences in means over time for a single group, a one-way repeated measures ANOVA with Geisser-Greenhouse correction was used. To analyze differences in means over time for multiple groups, a two-way repeated measures ANOVA with Geisser-Greenhouse correction was used. For both repeated measures ANOVA tests, post-hoc Dunnett’s tests were performed only in cases where there was a significant effect of time (one-way) or a significant interaction (two-way). Significance level α was set at 0.05, and when necessary P values were adjusted for multiple comparisons prior to comparing to α. All P values are reported in figure legends.

## Acknowledgments

We thank Julia Deere, Lisandro Martin, Jocelyn Yao, Nathan Liang, and Kerlina Liu for technical assistance; Arnold Heynen, Kiki Chu, Erin Hickey, Athene Wilson-Glover, and Jessica Buckey for research and administrative support; Victoria Mulloy, Victoria Donovan, and Chris Harvey-Clark for animal and veterinary support; Jeffrey Gavornik, Nathan Crowder, and Braden Kamermans for software development; Donald Mitchell and Lara Pierce for helpful discussions; and Eric Gaier for constructive comments on the manuscript. This work was supported by NIH K99 EY029326 to M-f.F., CIHR 153333 to K.R.D., and NIH R01 EY029245 to M.F.B.

## Supplemental Information

**Figure S1:**
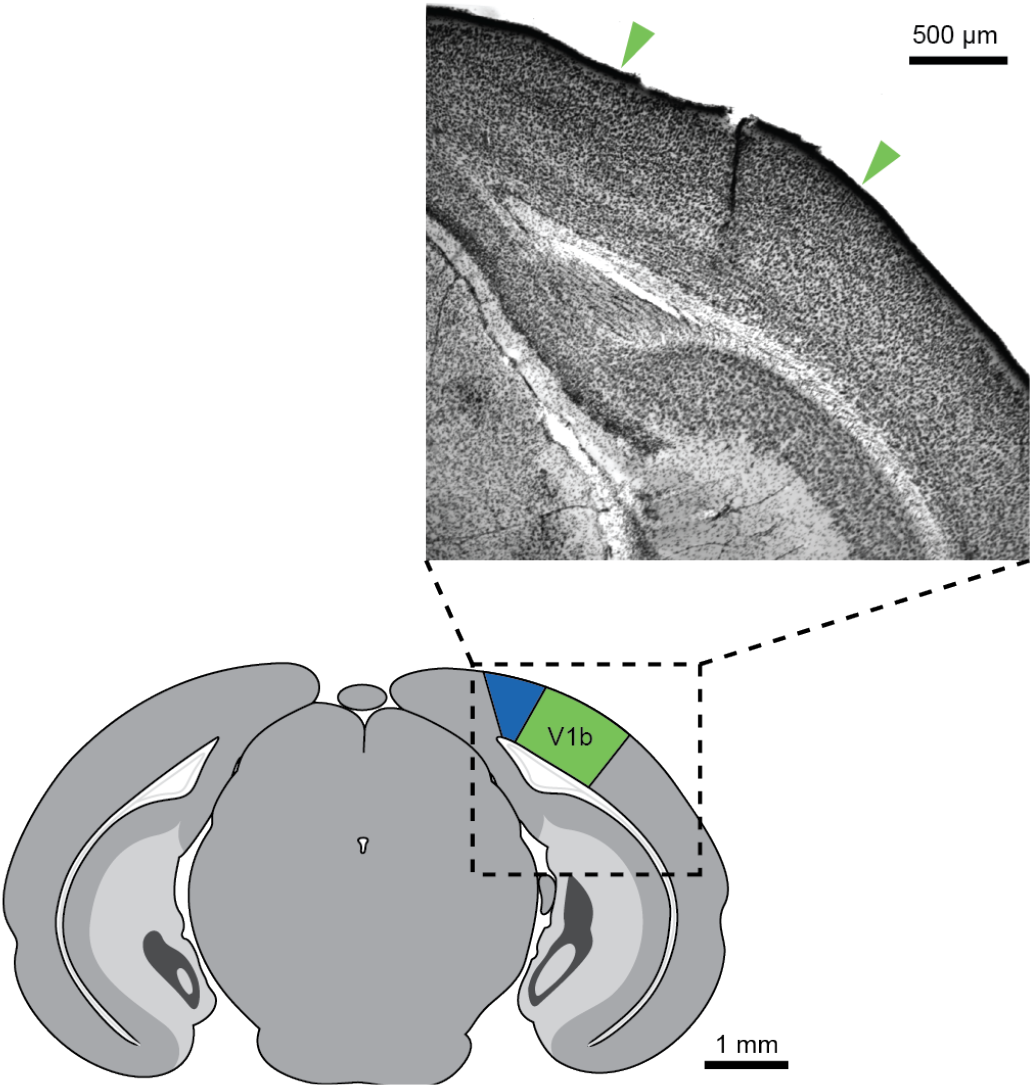
Histological verification of electrode position in mouse binocular V1. *Top*, Nissl-stained coronal slice of mouse brain showing track left by an electrode implanted in layer 4 of V1. Green arrows denote the boundaries of binocular V1. *Bottom*, cartoon of coronal slice from mouse brain corresponding to micrograph above. The monocular and binocular segments of V1 are shaded in blue and green, respectively.

**Figure S2:**
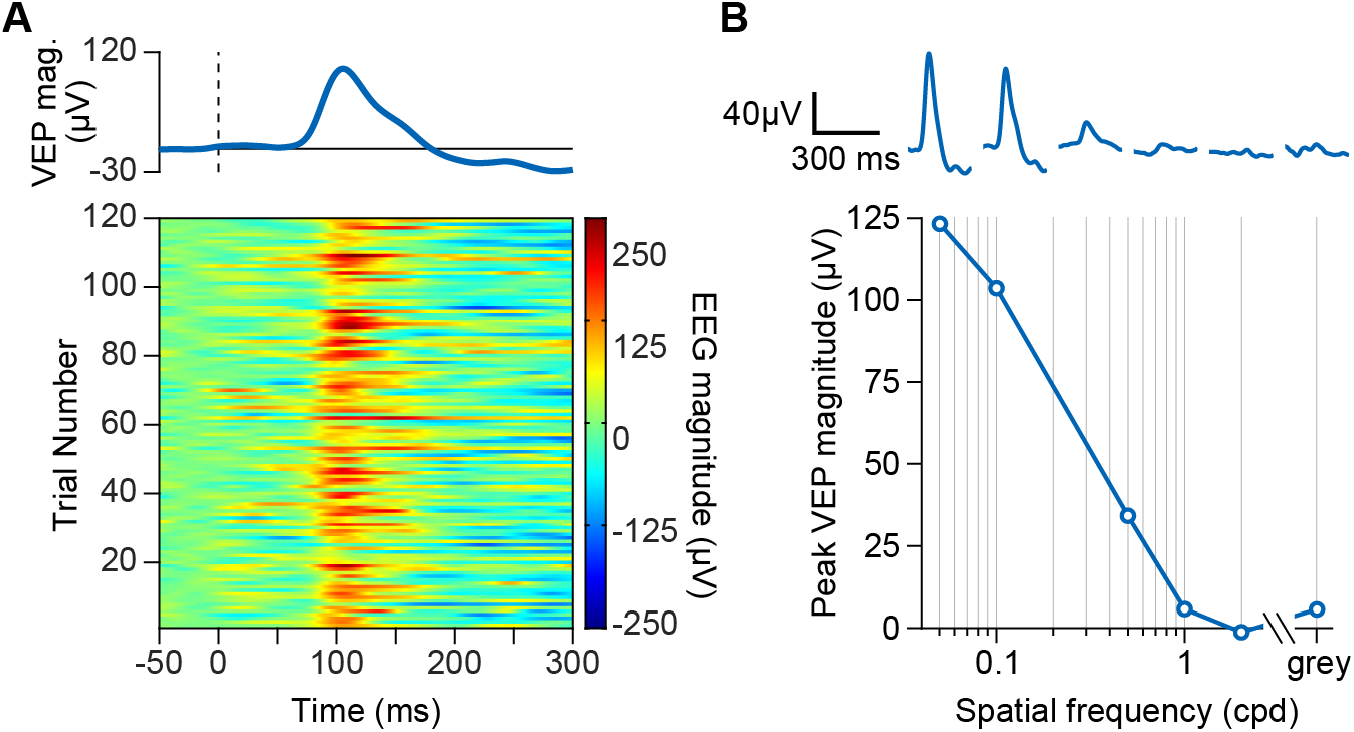
Time-domain VEPs from cat scalp surface field potential. **(A)** *Bottom*, example phase reversal-aligned EEG for 120 trials within a recording session where the 0.1 cpd grating stimulus was presented (interleaved with blocks of grating stimuli at other spatial frequencies). The EEG is normalized to a 50-millisecond window prior to each phase reversal at time = 0. *Top*, VEP computed as the average across all trials shown below. **(B)** *Top*, VEP waveforms for grating stimuli presented at (from left to right): 0.05 cpd, 0.1 cpd, 0.5 cpd, 1 cpd, 2 cpd, and grey screen. *Bottom*, peak VEP magnitude during the 300-ms window following the phase reversal, quantified as a function of spatial frequency. Data for 0.1 cpd is the same as shown in A, and data used in this plot is the same as example of frequency-domain analysis shown in Fig. 4C.

**Figure S3:**
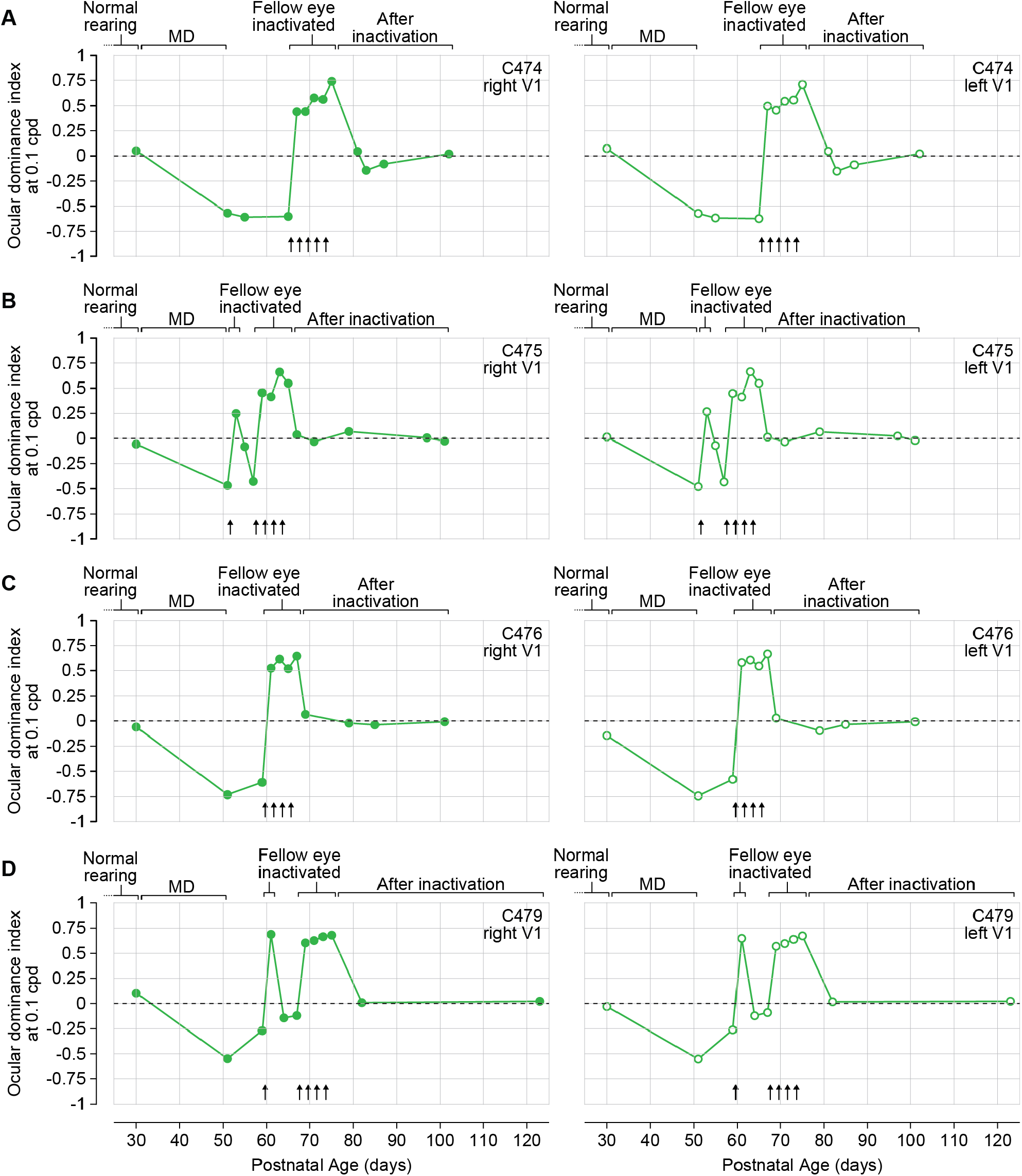
Trajectory of deprivation-driven ocular dominance shift and inactivation-mediated recovery in individual cats. Ocular dominance indices (ODIs) over time for 4 cats subjected to long-term MD followed fellow eye inactivation. ODIs are computed from scalp surface potential recordings during monocular viewing of grating stimuli at 0.1 cpd. Arrows denote time of TTX injections. For each plot, the dashed line at 0 indicates a balanced ODI typical of a visually normal animal, whereas values of 1 and −1 indicate complete dominance by either the left or right eye, respectively. Error bars, SEM.

**Table S1:** Monocular responses in mouse V1 after long-term MD (or sham) followed by fellow eye TTX (or saline)

**(A) Deprived contralateral eye**
| Group | *n* | Spatial freq. (cpd) | VEP magnitude (uV; **mean**± SEM) |  |  |  |  |
| --- | --- | --- | --- | --- | --- | --- | --- |
|  |  |  | P47<sup>‡</sup> | P54 | P61 | P68 | P75<sup>‡</sup> |
| Sham | 16 | 0.05 | **185.8** ± 17.20 | **173.34** ± 18.92 | **186.7** ± 20.08 | **164.9** ± 15.91 | **168.2** ± 15.95 |
|  |  | 0.2<sup>†</sup> | **195.1** ± 14.91 | **182.6** ± 13.12 | **190.5** ± 12.02 | **191.9** ± 15.70 | **194.0** ± 16.30 |
|  |  | 0.4 | **112.9** ± 13.54 | **111.3** ± 12.31 | **112.4** ± 11.94 | **101.8** ± 10.39 | **110.1** ± 9.690 |
| MD | 13 | 0.05 | **113.1** ± 13.90 | **129.6** ± 16.14 | **128.8** ± 13.99 | **96.57** ± 11.07 | **109.1** ± 16.70 |
|  |  | 0.2<sup>†</sup> | **105.2** ± 21.89 | **127.3** ± 18.30 | **132.7** ± 19.31 | **114.2** ± 14.83 | **116.1** ± 18.48 |
|  |  | 0.4 | **41.45** ± 7.197 | **66.54** ± 9.788 | **74.77** ± 13.31 | **64.08** ± 11.49 | **61.81** ± 13.60 |
| MD then TTX | 13 | 0.05 | **95.46** ± 13.05 | **156.5** ± 20.40 | **178.5** ± 21.87 | **166.6** ± 21.76 | **177.6** ± 25.15 |
|  |  | 0.2<sup>†</sup> | **95.84** ± 20.00 | **187.4** ± 25.83 | **214.5** ± 27.49 | **218.1** ± 23.39 | **210.8** ± 26.63 |
|  |  | 0.4 | **35.12** ± 7.723 | **93.13** ± 17.76 | **120.1** ± 19.95 | **108.3** ± 17.68 | **133.1** ± 26.37 |

<sup>†</sup>row represented in Fig. 2B-C; <sup>‡</sup>column represented in Fig. 2D

**(B) Fellow (non-deprived) ipsilateral eye**
| Group | *n* | Spatial freq. (cpd) | VEP magnitude (uV; **mean**± SEM) |  |  |  |  |
| --- | --- | --- | --- | --- | --- | --- | --- |
|  |  |  | P47<sup>‡</sup> | P54 | P61 | P68 | P75<sup>‡</sup> |
| Sham | 16 | 0.05 | **90.68** ± 10.17 | **89.55** ± 11.42 | **112.1** ± 18.39 | **83.90** ± 9.861 | **76.86** ± 8.851 |
|  |  | 0.2<sup>†</sup> | **91.36** ± 9.335 | **84.99** ± 9.324 | **105.6** ± 15.29 | **89.56** ± 9.759 | **74.67** ± 9.849 |
|  |  | 0.4 | **47.41** ± 7.675 | **45.58** ± 6.386 | **47.21** ± 8.373 | **43.83** ± 4.630 | **37.91** ± 5.341 |
| MD | 13 | 0.05 | **111.7** ± 20.39 | **75.77** ± 9.100 | **83.25** ± 16.79 | **62.41** ± 10.54 | **57.46** ± 9.788 |
|  |  | 0.2<sup>†</sup> | **118.717** ± 17.32 | **73.60** ± 10.30 | **85.13** ± 19.37 | **61.93** ± 9.419 | **64.09** ± 12.69 |
|  |  | 0.4 | **53.19** ± 9.23 | **40.82** ± 7.91 | **42.74** ± 9.553 | **31.64** ± 4.460 | **26.91** ± 4.213 |
| MD then TTX | 13 | 0.05 | **113.1** ± 17.38 | **123.4** ± 19.09 | **94.74** ± 14.03 | **79.69** ± 10.48 | **84.23** ± 14.57 |
|  |  | 0.2<sup>†</sup> | **130.2** ± 20.97 | **103.7** ± 21.61 | **102.7** ± 18.63 | **82.75** ± 15.47 | **83.52** ± 18.45 |
|  |  | 0.4 | **62.47** ± 14.99 | **49.47** ± 11.63 | **67.90** ± 14.17 | **40.96** ± 7.122 | **50.41** ± 11.37 |

<sup>†</sup>row represented in Fig. 2E-F; <sup>‡</sup>column represented in Fig. 2G

**Table S2:** Monocular responses in mouse V1 after long-term MD followed by fellow eye inactivation or reverse occlusion.

**(A) Deprived contralateral eye**
| Group | *n* | Spatial freq. (cpd) | VEP magnitude (uV; **mean**± SEM) |  |  |  |  |
| --- | --- | --- | --- | --- | --- | --- | --- |
|  |  |  | P47<sup>‡</sup> | P54<sup>‡</sup> | P61 | P68 | P75<sup>‡</sup> |
| MD then TTX | 14 | 0.05 | **95.40** ± 14.80 | **161.1** ± 21.79 | **146.7** ± 17.42 | **147.0** ± 16.20 | **170.1** ± 16.06 |
|  |  | 0.2<sup>†</sup> | **107.3** ± 18.19 | **171.3** ± 20.68 | **167.5** ± 19.29 | **164.1** ± 14.18 | **196.7** ± 16.32 |
|  |  | 0.4 | **42.59** ± 6.221 | **92.04** ± 14.85 | **107.1** ± 13.02 | **98.51** ± 11.47 | **109.6** ± 13.08 |
| MD then RO | 15 | 0.05 | **86.71** ± 8.53 | **154.0** ± 17.70 | **116.8** ± 14.99 | **95.24** ± 14.58 | **88.95** ± 14.30 |
|  |  | 0.2<sup>†</sup> | **104.4** ± 14.82 | **166.8** ± 12.44 | **128.5** ± 17.08 | **127.3** ± 20.00 | **97.30** ± 13.71 |
|  |  | 0.4 | **40.66** ± 6.209 | **91.49** ± 11.45 | **66.58** ± 11.02 | **63.22** ± 12.89 | **60.18** ± 9.32 |

<sup>†</sup>row represented in Fig. 3B-C; <sup>‡</sup>column represented in Fig. 3D

**(B) Fellow (non-deprived) ipsilateral eye**
| Group | *n* | Spatial freq. (cpd) | VEP magnitude (uV; **mean**± SEM) |  |  |  |  |
| --- | --- | --- | --- | --- | --- | --- | --- |
|  |  |  | P47<sup>‡</sup> | P54<sup>‡</sup> | P61 | P68 | P75<sup>‡</sup> |
| MD then TTX | 14 | 0.05 | **104.5** ± 9.991 | **93.29** ± 12.95 | **83.77** ± 13.28 | **75.40** ± 11.95 | **73.87** ± 9.574 |
|  |  | 0.2<sup>†</sup> | **119.3** ± 13.18 | **56.23** ± 5.962 | **58.09** ± 12.73 | **81.72** ± 19.80 | **60.89** ± 11.35 |
|  |  | 0.4 | **66.24** ± 9.288 | **30.58** ± 5.586 | **27.40** ± 9.281 | **42.14** ± 13.30 | **35.18** ± 5.402 |
| MD then RO | 15 | 0.05 | **96.99** ± 12.25 | **85.50** ± 22.25 | **71.12** ± 14.62 | **59.28** ± 14.51 | **41.92** ± 8.882 |
|  |  | 0.2<sup>†</sup> | **96.90** ± 11.60 | **52.57** ± 14.43 | **51.58** ± 10.62 | **38.81** ± 7.354 | **34.27** ± 7.008 |
|  |  | 0.4 | **56.77** ± 7.647 | **31.79** ± 4.858 | **24.52** ± 4.910 | **19.66** ± 4.196 | **22.75** ± 4.001 |

<sup>†</sup>row represented in Fig. 3E-F; <sup>‡</sup>column represented in Fig. 3G

**Table S3:** Ocular dominance indices in cat V1 after long-term MD followed by fellow eye TTX across spatial frequencies.

**(A) C474 (Top, right V1; Bottom, left V1)**
| Spatial freq. | Temporal frequency | Ocular dominance index |  |  |  |  |  |  |  |  |  |  |  |  |
| --- | --- | --- | --- | --- | --- | --- | --- | --- | --- | --- | --- | --- | --- | --- |
|  |  | P30<sup>†</sup> | P51<sup>‡</sup> | P55<sup>‡</sup> | P65<sup>‡</sup> | P67<sup>§</sup> | P69<sup>§</sup> | P71<sup>§</sup> | P73<sup>§</sup> | P75<sup>§</sup> | P81<sup>¶</sup> | P83<sup>¶</sup> | P87<sup>¶</sup> | P102<sup>¶</sup> |
| 0.05 cpd | 2 Hz | 0.064 | -0.617 | -0.497 | -0.460 | 0.522 | 0.511 | 0.592 | 0.516 | 0.686 | 0.182 | -0.061 | -0.034 | 0.030 |
|  | control | -0.077 | -0.123 | -0.029 | -0.044 | -0.015 | 0.054 | 0.079 | -0.063 | 0.073 | 0.022 | -0.102 | 0.081 | 0.004 |
| 0.1 cpd | 2 Hz<sup>#</sup> | 0.052 | -0.568 | -0.607 | -0.601 | 0.442 | 0.443 | 0.578 | 0.565 | 0.742 | 0.046 | -0.142 | -0.079 | 0.021 |
|  | control | -0.048 | -0.046 | -0.145 | -0.026 | -0.017 | 0.073 | 0.122 | 0.025 | -0.018 | -0.155 | 0.042 | 0.046 | -0.008 |
| 0.5 cpd | 2 Hz | -0.122 | -0.513 | -0.341 | -0.380 | 0.245 | 0.035 | 0.286 | 0.179 | 0.500 | 0.040 | -0.213 | -0.099 | 0.040 |
|  | control | 0.051 | -0.035 | -0.043 | 0.090 | -0.078 | -0.056 | 0.078 | 0.081 | 0.062 | 0.032 | -0.071 | 0.035 | 0.028 |
| grey | 2 Hz | 0.051 | -0.093 | -0.027 | 0.009 | 0.040 | 0.043 | -0.030 | -0.033 | 0.015 | 0.059 | -0.041 | -0.038 | -0.103 |
|  | control | -0.131 | -0.027 | -0.013 | -0.064 | 0.052 | 0.151 | -0.028 | 0.025 | 0.083 | -0.007 | -0.094 | 0.064 | 0.008 |
| 0.05 cpd | 2 Hz | 0.106 | -0.638 | -0.510 | -0.511 | 0.541 | 0.530 | 0.577 | 0.514 | 0.637 | 0.186 | -0.061 | -0.061 | 0.029 |
|  | control | -0.076 | -0.130 | -0.036 | -0.034 | -0.021 | 0.054 | 0.067 | -0.080 | 0.071 | 0.021 | -0.097 | 0.070 | 0.015 |
| 0.1 cpd | 2 Hz<sup>#</sup> | 0.074 | -0.571 | -0.616 | -0.622 | 0.497 | 0.455 | 0.545 | 0.558 | 0.710 | 0.047 | -0.151 | -0.088 | 0.022 |
|  | control | -0.019 | -0.036 | -0.131 | -0.038 | -0.020 | 0.033 | 0.123 | 0.014 | -0.069 | -0.138 | 0.040 | 0.104 | 0.019 |
| 0.5 cpd | 2 Hz | -0.151 | -0.485 | -0.366 | -0.381 | 0.283 | 0.022 | 0.259 | 0.194 | 0.477 | 0.045 | -0.206 | -0.096 | 0.042 |
|  | control | 0.026 | -0.023 | -0.044 | 0.061 | -0.085 | -0.039 | 0.058 | 0.091 | 0.053 | 0.048 | -0.070 | 0.022 | 0.011 |
| grey | 2 Hz | 0.039 | -0.073 | -0.028 | -0.016 | 0.053 | 0.071 | -0.045 | -0.029 | 0.070 | 0.002 | -0.051 | -0.090 | -0.102 |
|  | control | -0.123 | 0.017 | -0.010 | -0.040 | 0.031 | 0.127 | -0.015 | -0.005 | 0.040 | 0.005 | -0.087 | 0.052 | -0.013 |

<sup>†</sup>before MD; <sup>‡</sup>after MD; <sup>§</sup>during fellow eye inactivation; <sup>¶</sup>after fellow eye inactivation; <sup>#</sup>row represented in Figs. 4G and S3

**(B) C475 (Top, right V1; Bottom, left V1)**
| Spatial freq. | Temporal frequency | Ocular dominance index |  |  |  |  |  |  |  |  |  |  |  |  |  |
| --- | --- | --- | --- | --- | --- | --- | --- | --- | --- | --- | --- | --- | --- | --- | --- |
|  |  | P30<sup>†</sup> | P51<sup>‡</sup> | P53<sup>§</sup> | P55 | P57 | P59<sup>§</sup> | P61<sup>§</sup> | P63<sup>§</sup> | P65<sup>§</sup> | P67<sup>¶</sup> | P71<sup>¶</sup> | P79<sup>¶</sup> | P97<sup>¶</sup> | P101<sup>¶</sup> |
| 0.05 cpd | 2 Hz | -0.039 | -0.461 | 0.418 | 0.023 | -0.274 | 0.511 | 0.459 | 0.693 | 0.569 | 0.008 | -0.022 | 0.000 | -0.014 | 0.019 |
|  | control | -0.037 | 0.006 | 0.053 | 0.003 | -0.038 | -0.091 | -0.062 | -0.027 | -0.067 | 0.075 | 0.070 | 0.042 | 0.146 | 0.051 |
| 0.1 cpd | 2 Hz<sup>#</sup> | -0.055 | -0.467 | 0.250 | -0.081 | -0.428 | 0.457 | 0.416 | 0.661 | 0.550 | 0.037 | -0.031 | 0.069 | 0.007 | -0.025 |
|  | control | -0.074 | -0.045 | -0.064 | 0.086 | -0.104 | 0.080 | 0.067 | 0.165 | 0.117 | -0.052 | -0.014 | 0.036 | 0.033 | 0.008 |
| 0.5 cpd | 2 Hz | -0.131 | -0.393 | -0.017 | -0.154 | -0.555 | 0.263 | 0.124 | 0.274 | 0.186 | 0.179 | -0.158 | 0.205 | -0.043 | -0.062 |
|  | control | -0.006 | -0.051 | 0.007 | -0.002 | 0.050 | -0.044 | 0.017 | -0.044 | 0.034 | 0.055 | -0.070 | 0.063 | -0.061 | 0.072 |
| grey | 2 Hz | -0.074 | -0.026 | -0.038 | 0.044 | 0.058 | 0.001 | 0.020 | 0.055 | 0.107 | -0.059 | 0.097 | 0.015 | 0.039 | 0.021 |
|  | control | -0.124 | 0.020 | -0.008 | -0.033 | 0.029 | -0.047 | -0.075 | 0.149 | -0.055 | -0.003 | -0.031 | 0.062 | -0.078 | -0.061 |
| 0.05 cpd | 2 Hz | -0.056 | -0.469 | 0.383 | -0.040 | -0.297 | 0.515 | 0.455 | 0.699 | 0.569 | -0.011 | -0.020 | 0.003 | -0.013 | 0.029 |
|  | control | -0.033 | 0.025 | 0.028 | 0.003 | -0.052 | -0.052 | -0.024 | -0.008 | -0.067 | 0.068 | 0.072 | 0.035 | 0.076 | 0.021 |
| 0.1 cpd | 2 Hz<sup>#</sup> | 0.017 | -0.479 | 0.268 | -0.069 | -0.433 | 0.450 | 0.414 | 0.665 | 0.550 | 0.013 | -0.033 | 0.066 | 0.024 | -0.019 |
|  | control | -0.023 | -0.042 | -0.087 | 0.084 | -0.093 | 0.094 | 0.018 | 0.193 | 0.117 | -0.073 | 0.003 | 0.034 | 0.057 | -0.005 |
| 0.5 cpd | 2 Hz | -0.122 | -0.371 | -0.048 | -0.129 | -0.553 | 0.293 | 0.139 | 0.279 | 0.186 | 0.158 | -0.172 | 0.197 | -0.016 | -0.039 |
|  | control | -0.012 | -0.045 | 0.000 | -0.041 | 0.048 | -0.035 | 0.034 | -0.010 | 0.034 | 0.039 | -0.038 | 0.050 | -0.036 | 0.051 |
| grey | 2 Hz | -0.099 | -0.016 | -0.068 | 0.046 | 0.040 | 0.063 | 0.018 | 0.069 | 0.107 | -0.007 | 0.065 | 0.013 | 0.035 | 0.025 |
|  | control | -0.143 | 0.037 | -0.001 | -0.022 | 0.073 | -0.025 | -0.084 | 0.120 | -0.055 | -0.033 | -0.023 | 0.057 | -0.051 | -0.037 |

<sup>†</sup>before MD; <sup>‡</sup>after MD; <sup>§</sup>during fellow eye inactivation; <sup>¶</sup>after fellow eye inactivation; <sup>#</sup>row represented in Figs. 4G and S3

**(C) C476 (Top, right V1; Bottom, left V1)**
| Spatial freq. | Temporal frequency | Ocular dominance index |  |  |  |  |  |  |  |  |  |  |
| --- | --- | --- | --- | --- | --- | --- | --- | --- | --- | --- | --- | --- |
|  |  | P30<sup>†</sup> | P51<sup>‡</sup> | P59<sup>‡</sup> | P61<sup>§</sup> | P63<sup>§</sup> | P65<sup>§</sup> | P67<sup>§</sup> | P69<sup>¶</sup> | P79<sup>¶</sup> | P85<sup>¶</sup> | P101<sup>¶</sup> |
| 0.05 cpd | 2 Hz | 0.034 | -0.740 | -0.571 | 0.633 | 0.677 | 0.520 | 0.724 | 0.114 | -0.005 | 0.019 | 0.005 |
|  | control | -0.073 | -0.065 | 0.033 | 0.044 | 0.038 | -0.111 | 0.033 | 0.073 | 0.010 | -0.152 | 0.041 |
| 0.1 cpd | 2 Hz<sup>#</sup> | -0.058 | -0.733 | -0.610 | 0.525 | 0.617 | 0.519 | 0.646 | 0.066 | -0.021 | -0.038 | -0.008 |
|  | control | 0.009 | -0.092 | 0.078 | 0.047 | 0.025 | -0.022 | -0.036 | -0.009 | 0.205 | -0.032 | 0.035 |
| 0.5 cpd | 2 Hz | 0.027 | -0.700 | -0.537 | 0.381 | 0.410 | 0.390 | 0.382 | -0.001 | 0.144 | -0.100 | 0.016 |
|  | control | 0.039 | -0.028 | -0.053 | 0.031 | -0.049 | -0.098 | -0.008 | 0.053 | 0.099 | -0.023 | -0.040 |
| grey | 2 Hz | -0.070 | -0.002 | -0.059 | 0.114 | -0.078 | -0.074 | 0.089 | -0.019 | 0.175 | -0.094 | 0.052 |
|  | control | -0.005 | -0.147 | -0.073 | -0.012 | -0.053 | -0.046 | 0.145 | -0.103 | 0.133 | -0.087 | 0.032 |
| 0.05 cpd | 2 Hz | -0.070 | -0.754 | -0.560 | 0.657 | 0.649 | 0.552 | 0.728 | 0.077 | -0.049 | 0.026 | 0.010 |
|  | control | -0.037 | -0.067 | 0.051 | 0.054 | 0.013 | -0.091 | 0.018 | 0.075 | 0.036 | -0.156 | 0.032 |
| 0.1 cpd | 2 Hz<sup>#</sup> | -0.144 | -0.744 | -0.581 | 0.581 | 0.606 | 0.546 | 0.668 | 0.029 | -0.096 | -0.035 | -0.007 |
|  | control | 0.030 | -0.093 | 0.069 | 0.061 | 0.014 | -0.011 | -0.020 | 0.004 | 0.222 | -0.012 | 0.025 |
| 0.5 cpd | 2 Hz | 0.087 | -0.699 | -0.524 | 0.437 | 0.401 | 0.413 | 0.389 | -0.011 | 0.113 | -0.096 | 0.017 |
|  | control | 0.059 | -0.020 | -0.057 | 0.077 | -0.052 | -0.087 | -0.035 | 0.045 | 0.076 | -0.021 | -0.045 |
| grey | 2 Hz | -0.067 | -0.022 | -0.068 | 0.135 | -0.098 | -0.046 | 0.103 | -0.025 | 0.183 | -0.080 | 0.054 |
|  | control | 0.003 | -0.160 | -0.067 | 0.011 | -0.066 | -0.021 | 0.107 | -0.086 | 0.154 | -0.085 | 0.018 |

<sup>†</sup>before MD; <sup>‡</sup>after MD; <sup>§</sup>during fellow eye inactivation; <sup>¶</sup>after fellow eye inactivation; <sup>#</sup>row represented in Figs. 4G and S3

**(D) C479 (Top, right V1; Bottom, left V1)**
| Spatial freq. | Temporal frequency | Ocular dominance index |  |  |  |  |  |  |  |  |  |  |  |
| --- | --- | --- | --- | --- | --- | --- | --- | --- | --- | --- | --- | --- | --- |
|  |  | P30<sup>†</sup> | P51<sup>‡</sup> | P59 | P61<sup>§</sup> | P64 | P67 | P69<sup>§</sup> | P71<sup>§</sup> | P73<sup>§</sup> | P75<sup>§</sup> | P82<sup>¶</sup> | P123<sup>¶</sup> |
| 0.05 cpd | 2 Hz | -0.004 | -0.483 | -0.286 | 0.683 | -0.192 | -0.166 | 0.609 | 0.593 | 0.700 | 0.640 | -0.022 | 0.032 |
|  | control | 0.033 | 0.019 | 0.037 | -0.017 | 0.083 | 0.038 | -0.050 | -0.018 | -0.111 | 0.059 | 0.082 | 0.049 |
| 0.1 cpd | 2 Hz<sup>#</sup> | 0.103 | -0.547 | -0.269 | 0.689 | -0.143 | -0.119 | 0.604 | 0.627 | 0.666 | 0.681 | 0.009 | 0.023 |
|  | control | 0.005 | -0.050 | -0.007 | -0.030 | 0.063 | 0.000 | -0.078 | 0.072 | -0.044 | 0.026 | -0.063 | 0.054 |
| 0.5 cpd | 2 Hz | -0.046 | -0.451 | -0.386 | 0.510 | -0.362 | -0.290 | 0.487 | 0.329 | 0.451 | 0.499 | 0.033 | 0.037 |
|  | control | 0.047 | 0.001 | -0.018 | -0.027 | 0.070 | -0.040 | -0.069 | -0.049 | -0.021 | -0.004 | 0.019 | 0.054 |
| grey | 2 Hz | -0.074 | -0.043 | 0.024 | -0.007 | -0.008 | -0.041 | -0.048 | -0.011 | 0.148 | 0.126 | -0.121 | 0.098 |
|  | control | -0.129 | 0.108 | -0.001 | 0.018 | 0.010 | -0.012 | -0.066 | -0.062 | 0.003 | 0.069 | -0.039 | 0.060 |
| 0.05 cpd | 2 Hz | -0.058 | -0.497 | -0.278 | 0.647 | -0.136 | -0.123 | 0.588 | 0.550 | 0.652 | 0.643 | -0.033 | 0.034 |
|  | control | -0.030 | 0.017 | 0.007 | 0.867 | 0.091 | 0.022 | -0.085 | -0.015 | -0.081 | 0.066 | 0.072 | 0.045 |
| 0.1 cpd | 2 Hz<sup>#</sup> | -0.029 | -0.550 | -0.260 | 0.649 | -0.121 | -0.088 | 0.571 | 0.598 | 0.640 | 0.674 | 0.017 | 0.023 |
|  | control | -0.014 | -0.042 | -0.015 | 0.872 | 0.094 | 0.003 | -0.087 | 0.041 | -0.024 | 0.054 | -0.092 | 0.042 |
| 0.5 cpd | 2 Hz | -0.079 | -0.475 | -0.378 | 0.492 | -0.326 | -0.225 | 0.442 | 0.303 | 0.409 | 0.483 | 0.068 | 0.033 |
|  | control | 0.039 | -0.007 | -0.042 | 0.839 | 0.109 | 0.000 | -0.104 | -0.043 | 0.030 | -0.032 | 0.056 | 0.061 |
| grey | 2 Hz | -0.012 | -0.056 | -0.017 | -0.027 | 0.018 | -0.056 | -0.095 | -0.012 | 0.096 | 0.153 | -0.100 | 0.091 |
|  | control | -0.054 | 0.111 | 0.027 | 0.873 | 0.036 | -0.016 | -0.094 | -0.057 | -0.016 | 0.112 | -0.063 | 0.048 |

<sup>†</sup>before MD; <sup>‡</sup>after MD; <sup>§</sup>during fellow eye inactivation; <sup>¶</sup>after fellow eye inactivation; <sup>#</sup>row represented in Figs. 4G and S3

